# Enhancers mediate euchromatin hopping at chromatin contact points

**DOI:** 10.1101/2025.09.05.674255

**Authors:** Shanelle Mullany, Tiegh Taylor, Hangpeng Li, Yaqing Zhao, Alisha Suri, Tom Sexton, James O. J. Davies, Jennifer A. Mitchell

## Abstract

Enhancer-mediated gene activation involves the recruitment of chromatin modifiers and RNA polymerase to target promoters, but it is unknown if enhancers influence chromatin beyond their target genes. Euchromatin and heterochromatin associated histone modifications separate the genome into opposing nuclear compartments. Whereas heterochromatin marks are known to spread from one modified nucleosome to another, no such ability has been ascribed to euchromatin. Using mono-allelic enhancer deletions, native ChIP-seq, and an engineered interaction between an enhancer and transcriptionally inert DNA, we show that enhancers mediate the acquisition of euchromatin features at distal regions through chromatin looping. We term this phenomenon euchromatin hopping and found it occurring on average ∼270kb bidirectionally from enhancers, redefining our understanding of enhancer-mediated chromatin architecture with implications on enhancer identification using chromatin features.

**Graphical Abstract:** 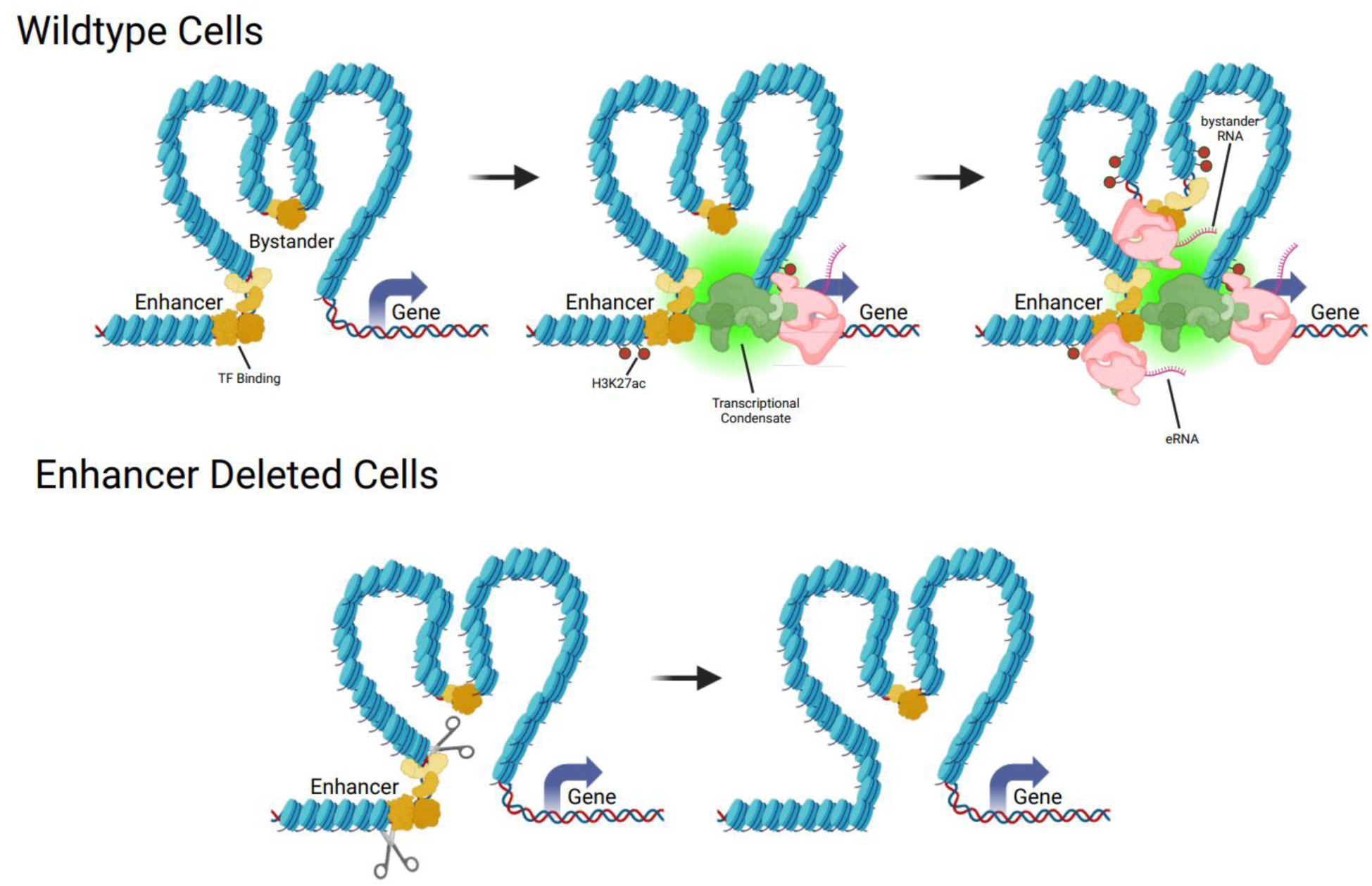

*Euchromatin hopping model:* Figure showing the proposed euchromatin hopping model. TFs recognize and bind to their binding sites in an active enhancer region. Upon activation enhancers recruit coactivators and RNAPII forming a condensate that supports gene activation. After an abundance of transcriptional machinery and coactivators are recruited, adjacent TF bound sites acquire euchromatin features through physical proximity to the active compartment, we call these regions “bystanders”. Upon enhancer deletion, condensate formation is lost and active euchromatin marks are not acquired at the gene promoter or other enhancer chromatin contacts. TFs are displayed in yellow, coactivators in green, RNAPII in pink, and histone modifications in red.

## Introduction

Enhancer elements are regions of non-coding DNA that positively regulate transcription of one or more genes to instruct complex processes like cell differentiation and tissue homeostasis ^1,2^. They regulate gene expression by binding multiple transcription factors (TFs), these TFs have intrinsically disordered protein domains ^3,4^ which aid in the recruitment of histone-modifying enzymes, co-activators, and RNA polymerase II (RNAPII) to a target gene ^5,6^. This cascade of interactions alters the chromatin landscape, where enhancers acquire specific histone modifications ^7^ and produce short non-coding enhancer RNAs (eRNAs)^8^, and genes gain an active transcriptional state^2^. H3K27 acetylation (ac) is a common histone modification found at enhancers ^9^ which is mutually exclusive to the H3K27 tri-methylation (me3) modification found in heterochromatin ^10^. Currently, enhancers are predicted by chromatin features; multiple TF binding, H3K27ac, H3K4 mono-methylation (me1), ^9,11,12^ and eRNA production ^9,12^, however these predictions have a high false positive rate (∼39-74%), revealed by massively parallel reporter assays ^13–17^. This indicates that many regions in the genome have enhancer-like features but lack independent regulatory activity. Through investigating specific enhancer predictions made by ENCODE ^12^ and FANTOM ^18^, we address this issue by determining how non-regulatory sequences with enhancer-like chromatin features emerge. We show that enhancers drive euchromatin hopping by imposing euchromatin features at chromatin contact regions, not just gene promoters but at many other loci within their topological domain. Understanding the broader genomic impact of enhancer activity is crucial for decoding the complex relationship between chromatin dynamics and gene activation across cell types.

## Results

### Enhancer deletion disrupts the epigenetic landscape and non-coding RNA production at surrounding euchromatin

To best study the relationship between enhancers and their surrounding chromatin landscape we used a hybrid cell line of mouse embryonic stem cells [mESC; 129/Sv:CAST/EiJ], allowing allele specific deletion, expression, and chromatin analysis due to the prevalence of single nucleotide polymorphisms (SNPs, ∼1/150bp). We focused on the locus of the *Sall1* gene, which encodes a TF dispensable in mESC, and to find the enhancers responsible for its regulation, we scanned for Multiple TF-bound Loci (MTL >= 4 TFs within 3.5kb) and found 4 downstream of the *Sall1* gene. Each MTL (28, 40, 52, and 92), labeled based on the distance in kb from the *Sall1* promoter, displays H3K27ac and produces eRNA-like nascent RNA ^19^ (Fig. 1A). Consistent with these chromatin features each region is a predicted enhancer in mESC by ENCODE3 ^12^, FANTOM ^18^, and ChromHMM ^20^ (Fig. 1A). MTL deletions (Table S1) followed by gene expression analysis, however, show only MTL28 and 40 display any regulatory activity in mESCs ^21^, together accounting for 98.6% of *Sall1* expression (Fig. 1B). By contrast, deletion of MTL52 or 92 does not significantly affect *Sall1* expression when deleted alone or in combination with the 2 active enhancers (MTL28/40)(Fig. 1B). Our previous study conducted an RNA-seq of cells with a deletion of MTL28-52 and saw *Sall1* was the only gene affected, indicating MTL52 is not regulating any other genes ^22^, and RNA expression assessed with Reverse Transcription Quantitative PCR (RT-qPCR) revealed MTL92 deletion does not influence the expression of other genes within its Topologically Associated Domain (TAD) (Fig. S1).

**Fig. 1.**
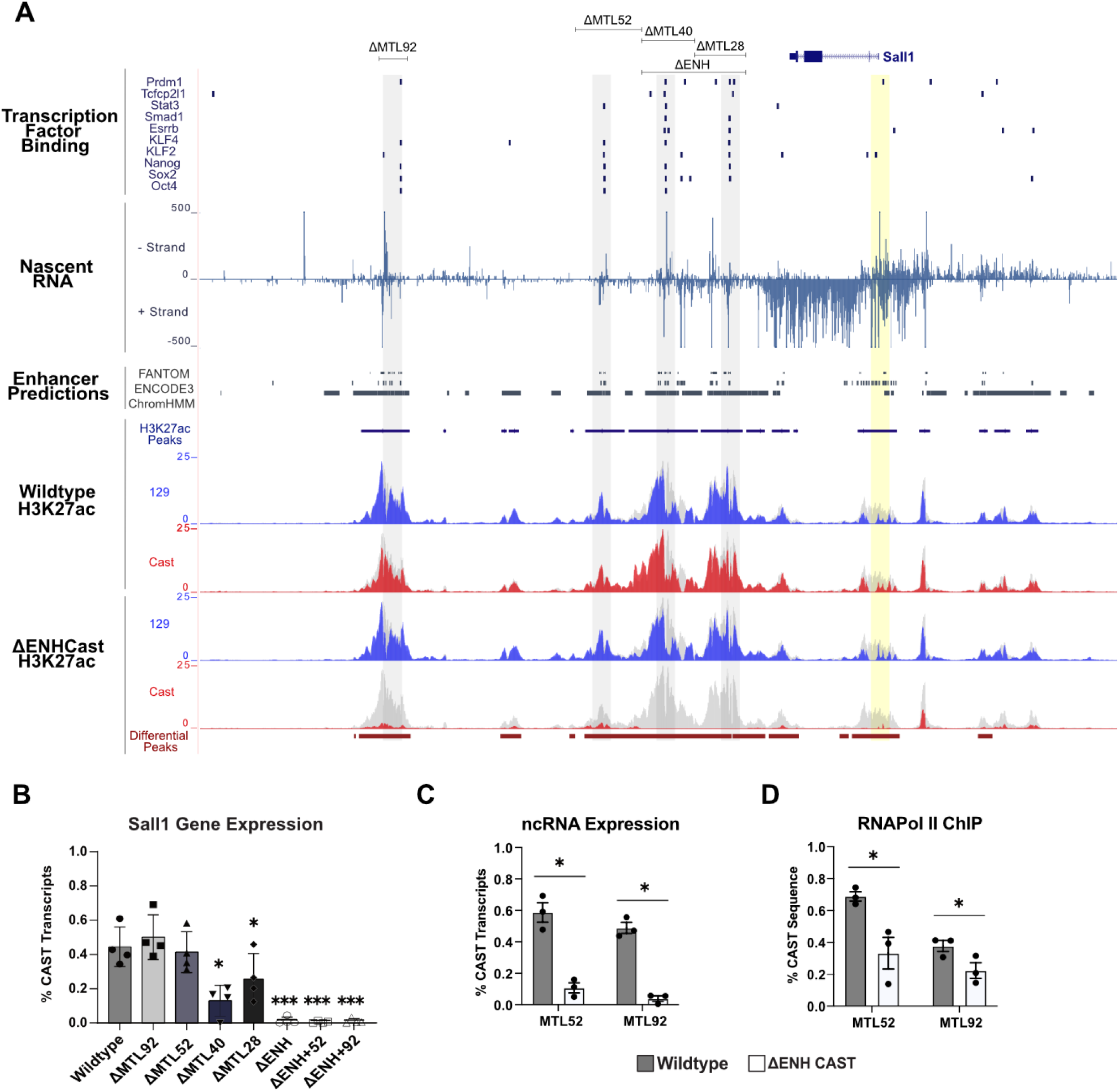
Active enhancers mediate chromatin feature propagation to local TF bound regions. (**A)** Multiple Transcription factor bound Loci (MTLs) near Sall1 are highlighted in grey along with their deletion sites marked by a black line at the top. Below are tracks for transcription factor binding ChIP data from CODEX data, bidirectional nascent RNA (Gro-Seq: GSE80262), and enhancer predictions from FANTOM, ENCODE 3, and ChromHMM. The Sall1 promoter is highlighted in yellow. Native H3K27ac ChIP-Seq data from wildtype and ΔENHCAST cells displaying reads from 129 in blue, CAST in red, overlaid on Total reads in grey. Peaks called from all reads are displayed in dark blue at the top and differential peaks between the wildtype and ΔENHCAST CAST allele in dark red at the bottom, all tracks displayed are in reads per million. **(B)** Sall1 gene expression of wildtype cells, each MTL deletion on a wildtype background, and each MTL deletion on the enhancer deleted background measured by RT-qPCR displaying % CAST transcripts (CAST transcripts/ 129 + CAST transcripts) * = p< 0.05, *** = p< 0.01. **(C)** ncRNA measured with RT-qPCR, grey is wildtype and white is ΔENHCAST, * = p< 0.05. **(D)** RNAPol II ChIP qPCR on wildtype in grey, and ΔENHCAST cells in white. % CAST sequence was calculated by (CAST sequence / (129 + CAST sequence)), * = p< 0.05.

Since MTL52 and 92 display enhancer-like features but lack independent regulatory activity in mESCs, we evaluated if the acquisition of these features depends upon the active enhancers MTL28 and 40. To execute this, we deleted both active enhancers (ΔENH; MTL28-40) on the CAST/EiJ (CAST) allele and surveyed chromatin changes within the locus. As the MTL deletions are mono-allelic we can evaluate changes to the chromatin on the deleted allele in comparison to the intact allele in the same *trans*-regulatory environment. In wildtype cells native H3K27ac ChIP-seq, performed without cross linking, showed almost identical enrichment on both the 129/Sv (129) and CAST alleles, with peaks at the four MTL and the *Sall1* promoter (Fig. 1A). Allele specific native H3K27ac ChIP-seq of clones with an MTL28-40 deletion on the CAST allele (ΔENHCAST, Fig. 1A) show H3K27ac enrichment was strikingly lost at the *Sall1* promoter (-4.35 fold; P= 0.0007), and the inactive MTL52 (-2.67 fold; P= 0.008) and MTL92 (- 4.45 fold; P= 0.0059). Throughout this locus several locations with weaker H3K27ac signal also display significant signal loss after enhancer deletion (Table S2), and each of these regions corresponds to locations bound by a few TFs. In contrast, the active enhancers are not dependent on MTL52 or 92 for H3K27ac enrichment as deletions of these MTL caused no change to H3K27ac at MTL28 or 40 (Fig. S2).

To assess the role of the MTL28 and 40 enhancers in the production of non-coding RNA (ncRNA) at MTL52 and 92 we conducted allele specific RT-qPCR in ΔENHCAST cells. In wildtype F1 mESC the distribution of ncRNA is close to equal between the 129 and CAST alleles at MTL52 and MTL92, with ∼60% of the transcripts originating from the CAST allele (Fig. 1C). Enhancer deletion caused a significant decrease in ncRNA production from the deleted allele at MTL52 (-5.48 fold, P= 0.0291) and MTL92 (-12.5 fold, P= 0.0132) (Fig. 1C). To determine if RNAPII recruitment was altered at these loci upon enhancer deletion, we performed allele specific ChIP-qPCR and observed a reduction in RNAPII recruitment to MTL52 (-1.94 fold, P= 0.0464) and MTL92 (-1.69 fold, P=0.0372) (Fig. 1D).

Conversely, ncRNA production at the enhancers was not dependent on MTL52 or 92 as their deletions had no effect on RNA levels at MTL28 or 40 (Fig. S3). Collectively, these findings indicate that H3K27ac, ncRNA production, and RNAPII recruitment at MTL52 and 92 are dependent on the active enhancers that drive *Sall1* transcription which cause enhancer-dependent euchromatin acquisition throughout the *Sall1* locus.

### Transcription factor binding is maintained at enhancer-dependent euchromatin

We considered the possibility that the active enhancers are supporting TF binding at MTL52 and 92, which could influence recruitment of histone modifiers and RNAPII. We focused on OCT4, SOX2, and KLF4 since these TFs can increase DNA accessibility, and recruit RNAPII and EP300 ^23–25^. Allele specific ChIP-seq showed evidence of OCT4, SOX2, and KLF4 binding to each of the 4 MTL on both alleles in wildtype cells (Fig. S4) and we identified strong matches to OCT4, SOX2 and KLF4 consensus motifs at each MTL (Table S3). To test if the binding of OCT4, SOX2, or KLF4, is altered at MTL52 or 92 after enhancer deletion we used allele specific ChIP-qPCR which revealed that KLF4 recruitment to MTL52 is significantly decreased (-1.60 fold, P = 0.0197), but not abolished (Fig. 2A). SOX2 and OCT4 also showed reduced recruitment to MTL52 after enhancer deletion, however, this difference was not significant. These findings suggest that the core TFs of the mESC gene regulatory network can access their TF binding site at the transcriptionally inert MTL even after enhancer deletion. In line with this, we found the enhancer deletion has no significant effect on chromatin accessibility, determined by ATAC-seq, at the *Sall1* promoter and MTL52 (Fig. S4). There was a significant decrease in chromatin accessibility at MTL92 (-5.48 fold, P= 0.0155), but this region did remain accessible. In contrast to the stark loss of H3K27ac and ncRNA after enhancer deletion, TFs remain capable of independently binding MTL52 and 92, and chromatin accessibility is largely maintained.

**Fig. 2.**
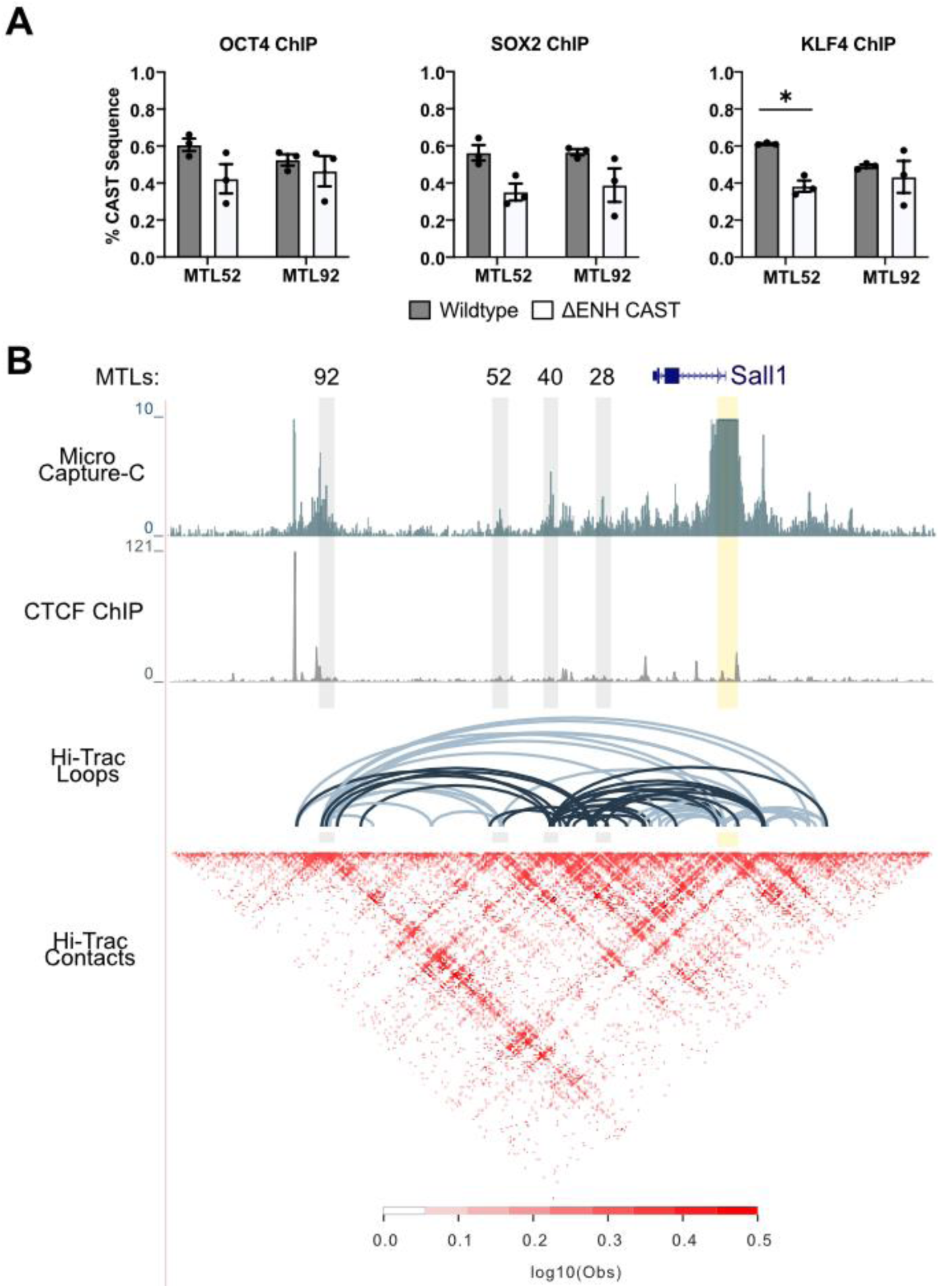
Euchromatin hopping occurs at TF bound regions displaying increased interaction with active enhancers. **(A)** ChIP-qPCR on wildtype in grey and ΔENHCAST cells in white, % CAST sequence was calculated by (CAST sequence / (129 + CAST sequence)), * = p< 0.05. **(B)** Micro Capture-C in wildtype E14 mESC with the Sall1 promoter as the viewpoint marked by a black triangle, and reads are shown in reads per million. Below is CTCF ChIP data (GSM699165) from the same cell type quantified by reads per million. Hi-Trac data is from E14TG2 showing chromatin contacts of open chromatin (GSE180175) displayed as called loops (Poisson P-value < 0.001), and an interaction matrix.

### Enhancer-dependent euchromatin is acquired at the chromatin contact regions of active enhancers

After finding enhancers MTL28 and 40 are influencing epigenetic modifications and ncRNA production at MTL52 and 92 without substantially altering TF binding, we considered spatial proximity as a mechanism for enhancer-mediated euchromatin feature acquisition. We analyzed Hi-Trac data from E14 mESC ^26^ and found significant chromatin interactions between the active enhancers (MTLs 28-40) and MTLs 52 and 92 (dark blue Hi-Trac Loops, Poisson P-value < 0.001, Fig. 2B), placing all 4 MTL in close three-dimensional space. To fine map these interactions we employed Micro Capture-C, which resolves chromatin-chromatin interactions at base-pair resolution ^27^, with the *Sall1* promoter as the viewpoint (Table S1). Strong interactions were observed between the promoter of *Sall1*, the active enhancers (MTLs 28, 40), as well as MTL52 and MTL92 (Fig. 2B). Although a strong interaction is captured between the *Sall1* promoter and a CTCF bound region, 5kb upstream from MTL92 (Fig. 2B), the MTL-promoter and MTL-MTL interactions occur at regions with minimal CTCF binding (Fig. 2B).

In summary, we show that the loss of H3K27ac, ncRNA, and RNAPII following enhancer deletion occurs at loci that physically interact with active enhancers. We propose that as enhancers recruit transcriptional machinery to activate their target promoters, nearby interacting regions acquire similar euchromatic features as bystanders. These observations led us to hypothesize that chromatin looping underlies this enhancer-driven euchromatin feature propagation, which we refer to as ’euchromatin hopping’.

### Engineering interaction with active enhancers induces euchromatin hopping

To investigate dependence on chromatin looping, we tested if an interaction between an active enhancer and a transcriptionally inert sequence is sufficient to cause euchromatin hopping. We used clones from a previous study ^28^ that inserted synthetic sequences between the *Sox2* gene and its distal enhancer that were engineered to contact the gene promoter (*Sox2* convergent CTCF motifs), enhancer (SCR convergent CTCF motifs), or neither region (Inert DNA) (Fig. 3 A-C). Importantly, these insertions had no effect on *Sox2* transcript levels, measured by RT-qPCR and RNA-FISH ^28^. Both CTCF insertions are 55 kb downstream of *Sox2* and contain the same sequence content but were inserted in different orientations, allowing us to determine if euchromatin-feature propagation is dependent upon sequence or enhancer contact. When surveying ncRNA levels with RT-qPCR at the insert site of each clone we found only the insert interacting with the active enhancer (SCR convergent) acquired any ncRNA (Fig. 3D). We then performed native H3K27ac ChIP-qPCR and again observed that only the insert interacting with the active enhancer (SCR Convergent) was able gain H3K27ac (Fig. 3E).

**Fig. 3.**
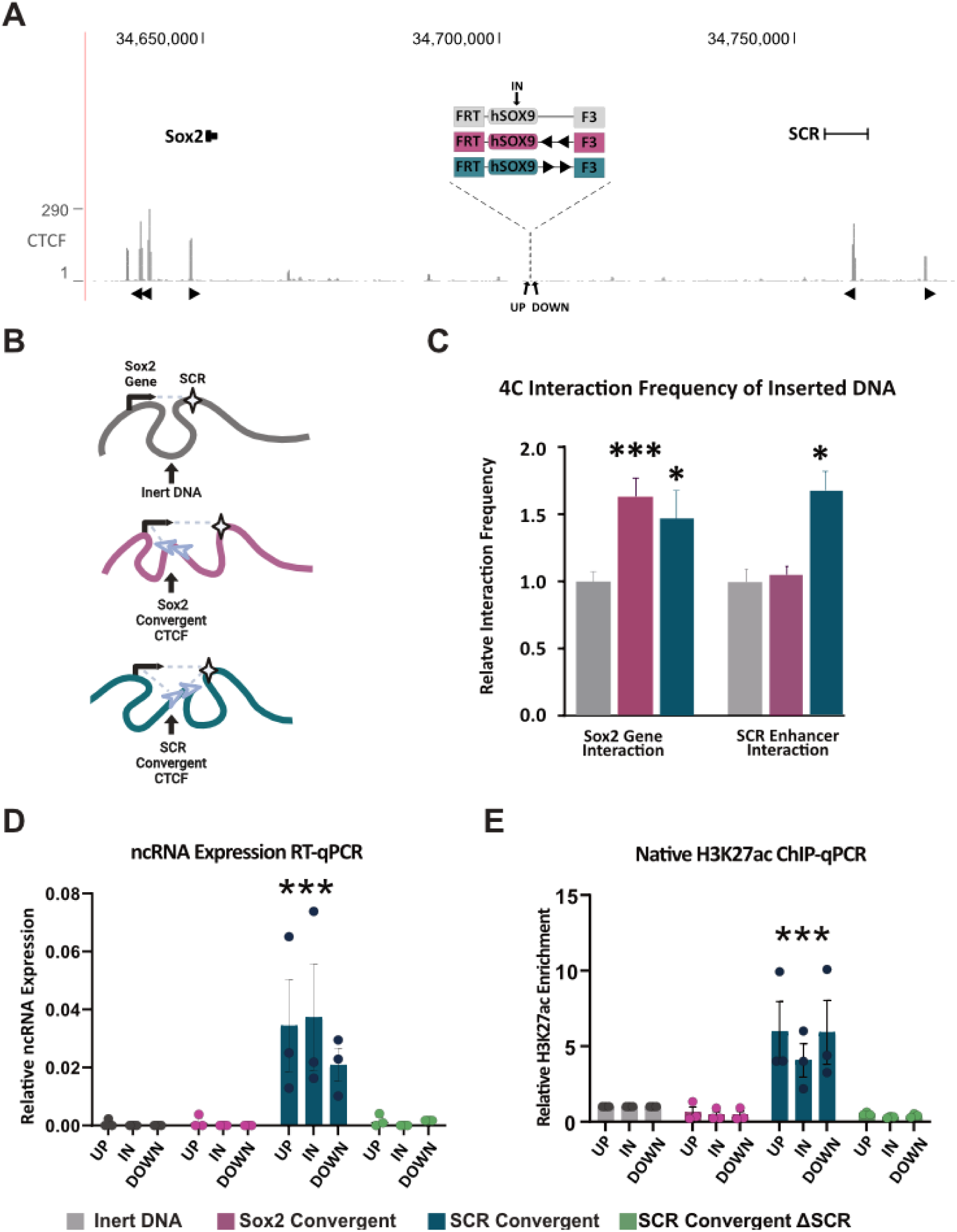
Engineering an interaction between an active enhancer and inert sequences causes euchromatin feature propagation. **(A)** The Sox2 locus with inserted sequences. CTCF sites and orientation shown as triangles, primer sites shown as arrows. **(B)** Schematic of the 3 different inserts and the chromatin architecture of each clone. SCR represents the Sox2 enhancer and triangles are the CTCF sites. **(C)** Quantification of interaction frequency between the insert and the Sox2 gene or Sox2 Enhancer (SCR) relative to the inert DNA insert was calculated using 4C data from these cell lines. **(D)** ncRNA expression at the insert site was calculated using RT-qPCR from each clone normalized to GAPDH expression. **(E)** Native H3K27ac ChIP-qPCR enrichment was normalized to a negative control and plotted relative to the inert DNA, * = P< 0.05, ***= P< 0.01.

The SCR is a super-enhancer known to form a transcriptional condensate enriched in cofactors such as RNAPII, Mediator, and histone acetyltransferases ^29–32^. Given this, we hypothesized that condensate formation by the SCR facilitates euchromatin hopping by delivering transcriptional and chromatin-modifying activities to bystander loci upon contact. Prior work demonstrated that deletion of the SCR disrupts condensate formation at the locus which reduces enhancer-promoter contacts and transcriptional bursting at *Sox2* ^30^. To test whether the condensate itself underlies euchromatin hopping, we deleted the SCR in the SCR-convergent clone and assessed both ncRNA production and H3K27ac at the insert. Upon SCR deletion, both ncRNA and H3K27ac enrichment at the insert site were lost (Fig. 3E), demonstrating that these features are dependent on the enhancer and its condensate-forming activity. Together, these results show that euchromatin hopping occurs specifically at bystander loci engaged in physical contact with an active enhancer, can activate transcriptionally inert loci, and may rely on the ability of an enhancer to form a condensate rich in transcriptional machinery.

### Euchromatin features propagate to adjacent accessible chromatin from allele specific enhancers genome-wide

To investigate if euchromatin hopping occurs genome-wide, we generated chromatin feature data from F1 mESCs (ATAC-seq, OCT4, SOX2, KLF4, H3K27ac, H3K4me1, and RNAPII ChIP-seq) and identified regions where strong enhancer features are exclusive to one allele. By identifying allele specific enhancers, we can determine if euchromatin hopping is occurring from these enhancers by surveying the surrounding chromatin on each allele. Variation in enhancer activity between alleles occurs due to SNPs that disrupt TF binding, resulting in differences to cofactor recruitment and overall enhancer function ^33,34^. To find allele specific enhancers we first identified the ∼40,000 differential ATAC-seq peaks between the 129 and CAST alleles (FDR <0.01) (34). At each differential ATAC peak, we calculated the fold change in read counts between the 129 and CAST alleles for ATAC, OCT4, SOX2, KLF4, H3K27ac, H3K4me1, and RNAPII. K-means clustering ^35^ of regions by fold change in allelic read counts identified four groups corresponding to strong and weak differential enrichment of all features (Fig. 4A). There are ∼4,000 regions with strong enrichment of all features on the 129 or CAST alleles. To determine to what extent these clusters may represent allele specific enhancers we used RNA-seq data from F1 mESCs to evaluate allele specific gene expression. We found most genes associated with the 2 strong differential enrichment clusters displayed allele specific gene expression that was significantly favoring the enriched allele and therefore classified these clusters as allele specific enhancers (Fig. S5). An example of a CAST enhancer is seen in Figure 4B (pink highlight), with CAST specific OCT4, SOX2, and KLF4 binding. Several ATAC peaks surrounding this enhancer show CAST-biased H3K27ac and H3K4me1, indicative of enhancer-driven euchromatin propagation similar to the *Sall1* locus (Fig. 4B, black arrowheads).

**Fig. 4.**
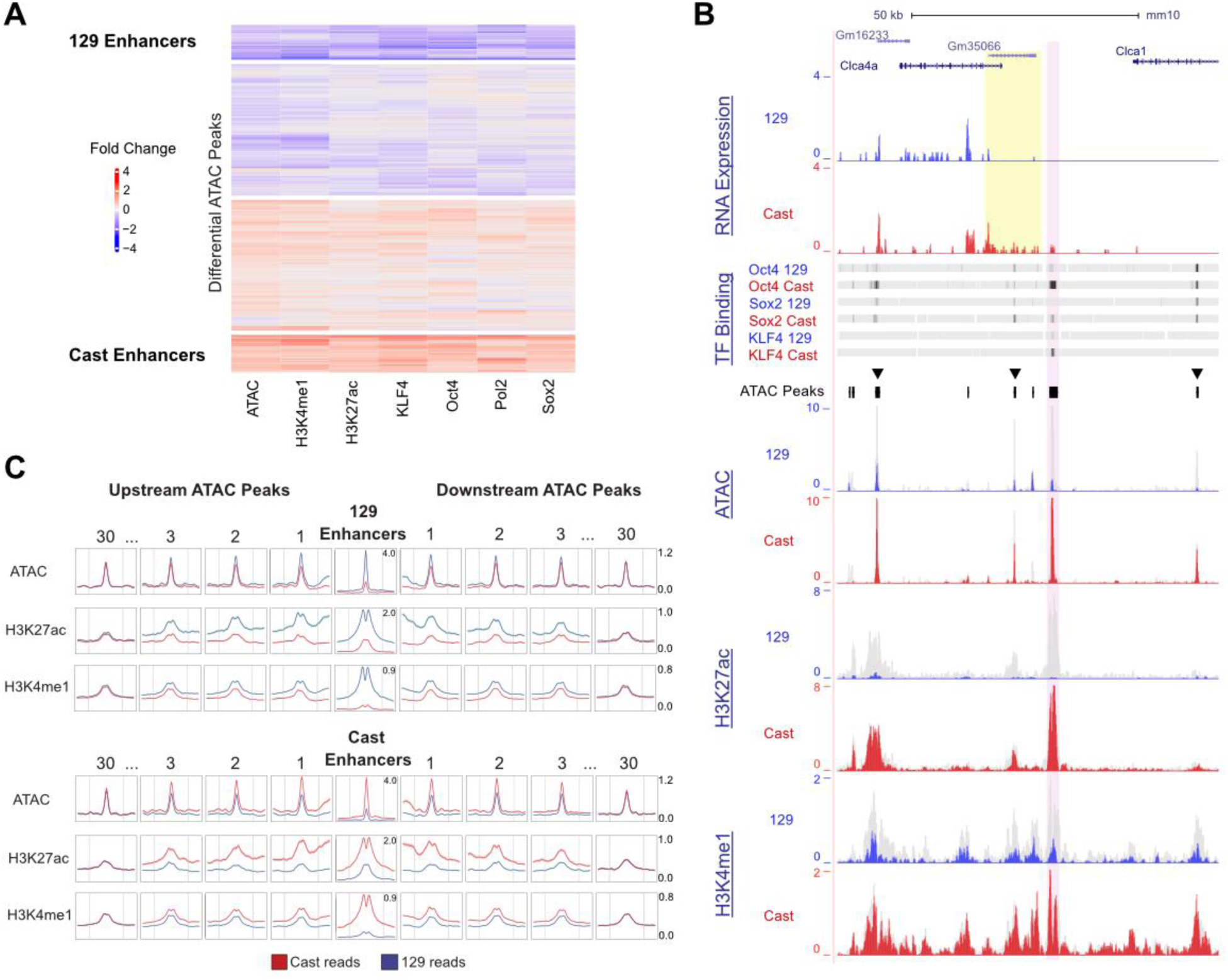
Euchromatin hopping occurs at allele biased enhancers throughout the genome. **(A)** A heatmap of all differential ATAC peaks and the fold change of read counts coming from 129 and CAST alleles calculated by (log2(CAST)) - (log2(129)), clustered by k-means clustering with Complex Heatmap. **(B)** A CAST specific enhancer is highlighted in pink, RNA from each allele is shown at the top and highlighted in yellow is a differentially expressed gene associated with the enhancer. Below is a heatmap of 129 and CAST binding events for Sox2, Oct4, and KLF4, calculated using ChIP-seq data and reads per million. Open chromatin peaks calculated from total reads are displayed underneath, followed by 129 and CAST bedgraphs of ATAC-Seq data shown in reads per million. Below is H3K27ac and H3K4me1 ChIP-seq data in reads per million. All 129 called reads are shown in blue, CAST in red, and total reads are displayed behind these trakcs in grey. Euchromatin hopping events are marked with a black arrowhead **(C)** NGS average profile plots of 129 and CAST reads for ATAC, H3K27ac, and H3K4me1 at ATAC peaks upstream and downstream of all 129 specific enhancers on top and CAST specific enhancers on the bottom. Average profile plots include a 4kb region centered on ATAC peaks and 1kb flank on either side. Peaks shown are the 1st, 2nd, 3rd, and 30th ATAC peak upstream and downstream from the enhancers.

To capture these events globally, we plotted the average profile of 129 and CAST enrichment for each feature at the allele specific enhancers. We then repeated this analysis at accessible chromatin peaks surrounding the enhancers, up to 40 peaks upstream and downstream (Fig. 4C, Table S4). We found the allelic read count significantly favored the allele containing the enhancer for ATAC, H3K27ac, H3K4me1 up to 21 peaks, upstream or downstream of the allele specific enhancer which is on average ∼270 kb away (Table S4, example peaks in Fig. 4C). This analysis suggests that enhancers are propagating euchromatin features, including H3K27ac and H3K4me1, to bystander regions at distances covering on average 540 kb. We also examined control regions that showed equal enrichment of enhancer-associated chromatin features on both the 129 and CAST alleles. For these biallelic enhancers, we assessed allelic read enrichment at nearby accessible chromatin peaks, up to 40 peaks upstream and downstream. Unlike the allele-specific enhancers, these biallelic regions showed no significant difference in 129 versus CAST read counts across the surrounding ATAC peaks (Fig. S6, Table S5), indicating that the allelic bias we observed surrounding allele specific enhancers is not due to read mapping or processing bias.

Since RNAPII accumulation in high abundance is associated with phase-separating, strong enhancers ^31,32,36,37^, we classified enhancers with ≥500 RNAPII reads per million as high RNAPII and those with lower levels as low RNAPII. We then analyzed the allelic enrichment of ATAC, H3K27ac, and H3K4me1 signal at the surrounding ATAC peaks for each group. We found that high RNAPII allele-specific enhancers were associated with euchromatin features extending further compared to low RNAPII enhancers (Fig. S7, Table S6). Specifically, in the high RNAPII group, open chromatin, H3K27ac, and H3K4me1 signals extended to approximately 9, 11, and 15 nearby peaks, respectively, compared to 4, 5, and 7 peaks in the low RNAPII group. Overall euchromatin features propagated ∼30kb further from the allele specific enhancers with high RNAPII signal compared to those with lower RNAPII enrichment (Table S6). These enhancers were also flanked by a greater number of accessible peaks, suggesting that stronger enhancer activity promotes euchromatin hopping to additional accessible regions. This supports the idea that transcriptionally active, phase-separating enhancers can propagate euchromatin features to surrounding chromatin.

## Discussion

Here we show enhancer mediated recruitment of transcriptional machinery is not exclusive to gene promoters but occurs at TF bound loci in three-dimensional proximity to active enhancers. These bystander regions acquire euchromatin features including histone modifications, accessibility, and ncRNA in an enhancer-dependent manner. Given that cell-type specific gene regulation relies on specific heterochromatin and euchromatin borders, euchromatin hopping might play a functional role in establishing and maintaining this landscape.

The ability of enhancers to induce euchromatin propagation aligns with the hub model of enhancer activity and their role in nuclear condensate formation ^38^. Many TFs have intrinsically disordered regions in their protein structure, and these domains have been found to phase separate, including CTCF ^39^, OCT4 ^3^, SOX2 ^40^, and KLF4 ^41^. Similar to what has been observed at the SCR ^29,30^, we propose the TFs bound at enhancers allow the formation of a compartment with an abundance of transcriptional machinery and coactivators ^39,42^. If this compartment contacts any neighbouring accessible sites bound by TFs with intrinsically disordered regions, euchromatin features are acquired and ncRNA production occurs in an enhancer-dependent manner, which appears as euchromatin hopping when mapped to a linear genome.

Given that histone modifications and nascent RNA are subject to enhancer-driven euchromatin hopping, and these features are used to predict the location of active enhancers ^12,43^, how can active enhancers be identified at genome scales with higher fidelity? We found that TF binding is less subject to euchromatin hopping, suggesting this feature may serve as a more robust predictor of enhancers ^44,45^. CRISPR deletion approaches ^21^, machine learning models ^46^, and synthetic enhancer assays ^47^ have revealed that enhancers typically contain multiple TF binding sites often exceeding ten per enhancer, whereas TF-bound regions with limited or no regulatory activity contain fewer TF binding sites. In support with this, the active MTL (28 and 40) at the *Sall1* locus are bound by 8-9 TFs and the inactive bystander MTL (52 and 92) are bound by only 6 TFs. Predicting the location and function of enhancers on a genome scale will require an understanding of where TFs are binding, and which TFs are functional together in a specific context or cell type.

Unlike heterochromatin spreading, which occurs linearly from one nucleosome to another, enhancer driven euchromatin hopping allows for the propagation of active chromatin features through chromatin contact points that can be several kb apart. This suggests that enhancer activity extends beyond direct transcriptional activation to include broader reconfiguration of the epigenetic landscape. Incorporating these insights into enhancer prediction models may provide a more accurate framework for deciphering regulatory elements across the genome.

## Resource Availability

The data from this study has been submitted to NCBI Gene Expression Omnibus (GEO; http://www.ncbi.nlm.nih.gov/geo/).

Series GSE247341; The following secure token has been created to allow review of record GSE247341 while it remains in private status: **qbijwycojxwtdan**

Series GSE247342; The following secure token has been created to allow review of record GSE247342 while it remains in private status: **qtirkaqyxfqxlyd**

Series GSE186864; The following secure token has been created to allow review of record GSE186864 while it remains in private status: **irknucqmxhczlib**.

## Supporting information

Supplemental Figues and Tables

## Acknowledgments

We thank the Mitchell lab for their input on this work, and the SickKids TCAG facility for all the high throughput sequencing. All funding came from Natural Sciences and Engineering Research Council of Canada 2020-05972 (JAM).

## Author contributions

Conceptualization: SM, TT, JAM

Methodology: SM, JD, TT

Investigation: SM, AS, HL, TS

Visualization: SM, TS, JAM

Funding acquisition: JAM

Writing- original draft: SM, JAM

Writing- review & editing: SM, TS, JAM

## Supplemental Information

Document S1. Figs. S1 to S7, Tables S1 to S6.

## STAR Methods

### Cell culture and CRISPR-Cas9 mediated monoallelic deletions

For all experiments F1 mESC (M. musculus129× M. CASTaneous), obtained from Barbara Panning, were used and grown on 0.1% gelatin coated 10cm plates in ES media comprised of DMEM, 15% FBS, 0.1 mM MEM nonessential amino-acids, 1 mM sodium pyruvate, 2 mM GlutaMAX, 0.1 mM 2-mercaptoethanol, 1000 U/mL LIF, 3 µM CHIR99021, and 1 µM PD0325901. CRISPR-Cas9 deletions were done using the protocol described in Moorthy & Mitchell 2016, Briefly, 5ug of Cas9-GFP, and 2 gRNAs were transfected using the Neon Transfection System into 1 million mESC. After 48-72 hours cells were taken to the FACS facility to be sorted for GFP and plated. These cells underwent colony picking and genotyping using allele specific primers to confirm the deletion.

### Chromatin Immunoprecipitation and sequencing

All cross-linked ChIP, done on Oct4, Sox2, KLF4, and RNAPII, were executed using a protocol adapted from Taylor et al 2022. In brief, cells were crosslinked with DSG for 50min and formaldehyde for 10min. The cells were then lysed, and chromatin was sonicated into 300bp fragments, where the sample was then split in 2. 500ul of sonicated chromatin was incubated with the appropriate antibody, anti-OCT4A (CST 5677), anti-SOX2 (CST 23064), anti-RNA polymerase II (Abcam ab5131), or anti-KLF4 (AF3158). Dynabeads were added and incubated with the samples overnight at four degrees. The next day lysates were washed with RIPA buffer 5 times and TBS once. The chromatin was then eluted, de-crosslinked, and treated with proteinase K before phenol chloroform precipitation. Allele specific primers were used to measure target enrichment via qPCR.

Native ChIP protocol was adapted from Nagano et al 2008, in summary, the nuclei of 10 million cells were isolated with a sucrose gradient. Following nuclei purification, the chromatin was fragmented to ∼150 bp using 15U of micrococcal nuclease in 500 ul volume for 3 minutes. The cells were then lysed and chromatin was incubated with the appropriate antibody overnight at 4 degrees, anti-H3K27ac (ab4729) and H3K4me1 (ab176877). The next day 60ul of protein A and protein G Dynabeads were added and incubated with antibody-bound chromatin overnight at 4 degrees. The next day samples were washed 3 times each with low, medium, and high salt buffers, before being eluted and incubated with proteinase K for 3 hours. Phenol chloroform precipitation was done to purify all samples before allele specific qPCR and sequencing.

Allele specific qPCR was performed using primers that carried a SNP between the 129 and CAST allele in the 3’ end. Expression values were calculated using a standard curve determined by serial dilutions of input samples that were not immunoprecipitated. In genome-wide sequencing experiments samples were sent to TCAG for library prep and sequenced using NovaSeq 6000 with 150bp paired-end reads at a depth of ∼50 million reads/sample. Sequencing data were trimmed using fastp and aligned using Bowtie2 to a custom genome with N-masking at all discriminatory SNPs between the 129 and CAST allele. Following alignment, reads were allele-called using the SNPsplit program. Duplicates and mitochondrial reads were removed with Samtools and Picard. Wildtype replicates prior to allele calling were used to call peaks using the Genrich tool, and DEseq2 was used to calculate differential peaks. Finally, bedgraphs were made using the bamCoverage tool.

### RNA isolation and expression analysis via RT-qPCR

RNA isolation and expression analysis protocol was carried out as described in Moorthy et al.^22^ where RNA was extracted from confluent 10cm plates using the RNeasy plus mini kit (Qiagen). An additional 30 minute incubation with turbo DNase was executed. RNA was reverse transcribed using the Multiscribe Reverse Transcriptase (Thermo Fisher). Samples were diluted to 200ng/ul stocks and expression was measured using qPCR with allele specific primers. A standard curve was generated using genomic DNA to normalize for different primer efficiencies. The allelic ratio of expression in each clone was calculated (CAST expression / (129 expression + CAST expression)). Significant differences were determined using GraphPads paired t-test calculator.

### Measuring genome wide chromatin accessibility with Assay for Transposase Accessible Chromatin with Sequencing (ATAC-Seq)

ATAC-Seq was executing following the improved Omni-ATAC protocol described by Cocres et al (Corces et al., 2017). Briefly, 60,000 cells were mixed with 50ul of Resuspension Buffer with final concentrations of reagents being: 0.1% NP40, 0.1% Tween-20, and 0.01% Digitonin. After 3 minutes of incubation on ice cells were lysed with 1mL Tween buffer composed of 0.1% Tween-20. Nuclei were then pelleted and resuspended in 50ul master mix supplemented with transposase, and reaction was incubated at 37 degrees for 30 minutes. Transposed fragments were cleaned up with MinElute columns (Qiagen). The libraries were prepared using the NEBNext HiFi kit, size selected, and sequenced with NovaSeq 6000 using paired- end 50 cycle with 50-60 million reads per sample. Sequencing data was analyzed using the same methods as Native ChIP.

### Identifying chromatin contacts using Micro Capture-C and Hi-TrAC data

Micro Capture-C was executed following the protocol described by Hua et al. ^27^. Briefly, for each sample 10 million cells were fixed with 2% final volume of formaldehyde in 10 ml media for 10 minutes. Cross-linked samples were then permeabilized with 1% digitonin and snap frozen in a mixture of dry ice and ethanol. Chromatin was then digested using micrococcal nuclease for 1 hour, leaving mostly mononucleosomes with linker DNA. Digested chromatin was then ligated overnight with T4 Polynucleotide Kinase PNK, DNA Polymerase I Large (Klenow) Fragment and T4 ligase (Thermo Fisher) and purified using the Qiagen DNeasy kit. 7-10 µg of each 3C library was sonicated to an average fragment size of 200 bp using a Covaris S220 Focused Ultrasonicator. The sample was then cleaned using Ampure XP beads. Libraries were prepared using the NEBNext kit and Ultra II ligation master mix. Following library prep, biotinylated oligonucleotide probe was used to capture the viewpoint. 10-20 µg of each library were combined and hybridization reactions were undertaken using Roche HyperCapture Target Enrichment Kit reagents (Roche, cat. no. 9075828001) following a modified protocol ^48^. Samples were then purified without size-selection using the AMPure XP beads. Sequencing was performed on the Illumina platform with 300 bp reads, utilizing paired-end 150 bp reads.

For Micro Capture-C analysis, adapters were trimmed using TrimGalore, and overlapping reads were merged into single sequences with FLASH by reconstructing fragments. The reconstructed fragments were then mapped to the oligonucleotide DNA sequence ±400 bp using BLAT to identify ligation junctions. This enabled the division of reads into new paired FASTQ files using MCCsplitter.pl, followed by alignment to the mouse reference genome mm10 with Bowtie 2. PCR duplicates were removed from the alignment files using MCCanalyser.pl. Both MCCsplitter.pl and MCCanalyser.pl are available for academic use via the Oxford University Innovation software store (https://process.innovation.ox.ac.uk/software/p/16529a/micro-capture-c-academic/1).

Hi-TrAC data (GSE180175) was input into cLoops2 and the script was followed as recommended on the github: https://github.com/YaqiangCao/cLoops2/. To call significant loops between MTLS parameters were set to show only 10kb or greater interactions that occurred minimum of 5 times; cLoops2 callLoops -d $chr -o $chr -eps 500 -minPts 15 -cut 10000 - max_cut -w -j.

### Engineered Interaction Between Inert DNA and Sox2 Enhancer

To engineer an interaction between inert DNA and the SCR a cassette with FRT and F3 sites were added 55kb away from the Sox2 promoter as described in Taylor et al., 2022.

Recombination allowed for the insert of CTCF sites in either orientation, which included the inert DNA as well. The inert DNA sequence was used to design primers and were the targets of each qPCR along with genomic regions flanking the insert. Interaction frequencies were determined using these coordinates for Sox2 (mm10; chr3: 34,644,922-34,664,967) and these for SCR (mm10; chr3: 34,749,652-34,760,919).

### Genome Wide Analysis of Allele Specific Enhancers

Differential ATAC peaks were called between 129 and CAST alleles using Diffbind (P<0.05). These differential peaks were adjusted to be 4kb and regions overlapping transcription start sites were removed. These regions were input into multibamsummary to get allele specific read counts of OCT4, Sox2, KLF4, RNAPII, H3K27ac, H3K4me1, and ATAC from allele specific bam files derived from SNPsplit. Read counts were turned into fold change between 129 and CAST alleles and the matrix was input into RStudio. ComplexHeatmap with K-means clustering of 4 clusters revealed strong and weak enriched sites for each allele. The strongest clusters were put into Genomic Region Enrichment Annotation Tool (GREAT) ^49^ along with a list of expressed genes for association. Regions associated with a gene outside of it’s TAD were filtered out and Enhanced Volcano was used to visualize gene expression of all genes associated with 129 and CAST specific enhancers. For each allele specific enhancer region, the nearest 40 ATAC peaks upstream and downstream were found and the BED coordinates were used in NGSplot to get average profile plots. Statistical significant read counts between 129 and CAST alleles were determined using a t-test, and the p-value was adjusted for sample size.

### Quantifying Allele Specific Enhancer Regulatory Capacity

To establish an integrated reporter cell line we modified our F1 mESC using CRISPR-Cas9 and this gRNA “GTGAGGGCTGGACTGCGAAC” to tag the Sox2 gene with Blue Fluorescent Protein (BFP) on the 129 allele and the mStrawberry fluorophore on the CAST allele. Once expression was confirmed with FACS and genotyping, we integrated FRT and F3 recombination sites downstream of the tagged Sox2-BFP sequence using this guide “AAGTTTTCTAGTCGGCATCA”. In between these recombination sites is a sequence encoding infrared protein (miRFP) driven by the phosphoglycerate kinase 1 (PGK) promoter and integrations were confirmed with FACS and genotyping. The SCR was deleted from these clones and deletion was confirmed with genotyping and the loss of BFP expression determined by FACS.

To assess the regulatory capacity of the allele specific enhancer example we extracted the 129 or Cast sequence of the locus using primers with a SNP favouring the desired allele. These sequences were then blunt cloned into a plasmid in between FRT and F3 sites. This plasmid along with a plasmid encoding the Flippase enzyme^28^ was then transfected into 1 million integrated reporter cells at a 5ug concentration. After integration was confirmed, BFP was quantified using FACS and compared to the parent cell line and a cell line with the Sox2 enhancer inserted into the FRT/F3 site.

## References

1. Arnone, M.I., and Davidson, E.H. (1997). The hardwiring of development: organization and function of genomic regulatory systems. Development 124, 1851–1864. 10.1242/dev.124.10.1851.

2. Banerji, J., Rusconi, S., and Schaffner, W. (1981). Expression of a beta-globin gene is enhanced by remote SV40 DNA sequences. Cell 27, 299–308. 10.1016/0092-8674(81)90413-x.

3. Boija, A., Klein, I.A., Sabari, B.R., Dall’Agnese, A., Coffey, E.L., Zamudio, A.V., Li, C.H., Shrinivas, K., Manteiga, J.C., Hannett, N.M., et al. (2018). Transcription Factors Activate Genes through the Phase-Separation Capacity of Their Activation Domains. Cell 175, 1842–1855.e16. 10.1016/j.cell.2018.10.042.

4. Triezenberg, S.J. (1995). Structure and function of transcriptional activation domains. Curr. Opin. Genet. Dev. 5, 190–196. 10.1016/0959-437x(95)80007-7.

5. Dynan, W.S., and Tjian, R. (1983). The promoter-specific transcription factor Sp1 binds to upstream sequences in the SV40 early promoter. Cell 35, 79–87. 10.1016/0092-8674(83)90210-6.

6. Vieira, K.F., Levings, P.P., Hill, M.A., Crusselle, V.J., Kang, S.-H.L., Engel, J.D., and Bungert, J. (2004). Recruitment of transcription complexes to the beta-globin gene locus in vivo and in vitro. J. Biol. Chem. 279, 50350–50357. 10.1074/jbc.M408883200.

7. Jenuwein, T., and Allis, C.D. (2001). Translating the histone code. Science 293, 1074–1080. 10.1126/science.1063127.

8. Vanhille, L., Griffon, A., Maqbool, M.A., Zacarias-Cabeza, J., Dao, L.T.M., Fernandez, N., Ballester, B., Andrau, J.C., and Spicuglia, S. (2015). High-throughput and quantitative assessment of enhancer activity in mammals by CapStarr-seq. Nat. Commun. 6, 6905. 10.1038/ncomms7905.

9. Creyghton, M.P., Cheng, A.W., Welstead, G.G., Kooistra, T., Carey, B.W., Steine, E.J., Hanna, J., Lodato, M.A., Frampton, G.M., Sharp, P.A., et al. (2010). Histone H3K27ac separates active from poised enhancers and predicts developmental state. Proc. Natl. Acad. Sci. U. S. A. 107, 21931–21936. 10.1073/pnas.1016071107.

10. Pasini, D., Malatesta, M., Jung, H.R., Walfridsson, J., Willer, A., Olsson, L., Skotte, J., Wutz, A., Porse, B., Jensen, O.N., et al. (2010). Characterization of an antagonistic switch between histone H3 lysine 27 methylation and acetylation in the transcriptional regulation of Polycomb group target genes. Nucleic Acids Res. 38, 4958–4969. 10.1093/nar/gkq244.

11. Heintzman, N.D., Hon, G.C., Hawkins, R.D., Kheradpour, P., Stark, A., Harp, L.F., Ye, Z., Lee, L.K., Stuart, R.K., Ching, C.W., et al. (2009). Histone modifications at human enhancers reflect global cell-type-specific gene expression. Nature 459, 108–112. 10.1038/nature07829.

12. Luo, Y., Hitz, B.C., Gabdank, I., Hilton, J.A., Kagda, M.S., Lam, B., Myers, Z., Sud, P., Jou, J., Lin, K., et al. (2020). New developments on the Encyclopedia of DNA Elements (ENCODE) data portal. Nucleic Acids Res. 48, D882–D889. 10.1093/nar/gkz1062.

13. Arnold, C.D., Gerlach, D., Stelzer, C., Boryń, Ł.M., Rath, M., and Stark, A. (2013). Genome-wide quantitative enhancer activity maps identified by STARR-seq. Science 339, 1074–1077. 10.1126/science.1232542.

14. Kwasnieski, J.C., Fiore, C., Chaudhari, H.G., and Cohen, B.A. (2014). High-throughput functional testing of ENCODE segmentation predictions. Genome Res. 24, 1595–1602. 10.1101/gr.173518.114.

15. Kheradpour, P., Ernst, J., Melnikov, A., Rogov, P., Wang, L., Zhang, X., Alston, J., Mikkelsen, T.S., and Kellis, M. (2013). Systematic dissection of regulatory motifs in 2000 predicted human enhancers using a massively parallel reporter assay. Genome Res. 23, 800– 811. 10.1101/gr.144899.112.

16. Zheng, Y., and VanDusen, N.J. (2023). Massively Parallel Reporter Assays for High-Throughput In Vivo Analysis of Cis-Regulatory Elements. J. Cardiovasc. Dev. Dis. 10, 144. 10.3390/jcdd10040144.

17. Barakat, T.S., Halbritter, F., Zhang, M., Rendeiro, A.F., Perenthaler, E., Bock, C., and Chambers, I. (2018). Functional Dissection of the Enhancer Repertoire in Human Embryonic Stem Cells. Cell Stem Cell 23, 276–288.e8. 10.1016/j.stem.2018.06.014.

18. Lizio, M., Abugessaisa, I., Noguchi, S., Kondo, A., Hasegawa, A., Hon, C.C., de Hoon, M., Severin, J., Oki, S., Hayashizaki, Y., et al. (2019). Update of the FANTOM web resource: expansion to provide additional transcriptome atlases. Nucleic Acids Res. 47, D752–D758. 10.1093/nar/gky1099.

19. Engreitz, J.M., Haines, J.E., Perez, E.M., Munson, G., Chen, J., Kane, M., McDonel, P.E., Guttman, M., and Lander, E.S. (2016). Local regulation of gene expression by lncRNA promoters, transcription and splicing. Nature 539, 452–455. 10.1038/nature20149.

20. Ernst, J., and Kellis, M. (2010). Discovery and characterization of chromatin states for systematic annotation of the human genome. Nat. Biotechnol. 28, 817–825. 10.1038/nbt.1662.

21. Singh, G., Mullany, S., Moorthy, S.D., Zhang, R., Mehdi, T., Tian, R., Duncan, A.G., Moses, A.M., and Mitchell, J.A. (2021). A flexible repertoire of transcription factor binding sites and a diversity threshold determines enhancer activity in embryonic stem cells. Genome Res. 31, 564–575. 10.1101/gr.272468.120.

22. Moorthy, S.D., Davidson, S., Shchuka, V.M., Singh, G., Malek-Gilani, N., Langroudi, L., Martchenko, A., So, V., Macpherson, N.N., and Mitchell, J.A. (2017). Enhancers and super-enhancers have an equivalent regulatory role in embryonic stem cells through regulation of single or multiple genes. Genome Res. 27, 246–258. 10.1101/gr.210930.116.

23. Ghaleb, A.M., and Yang, V.W. (2017). Krüppel-like factor 4 (KLF4): What we currently know. Gene 611, 27–37. 10.1016/j.gene.2017.02.025.

24. Mayran, A., and Drouin, J. (2018). Pioneer transcription factors shape the epigenetic landscape. J. Biol. Chem. 293, 13795–13804. 10.1074/jbc.R117.001232.

25. Shi, G., and Jin, Y. (2010). Role of Oct4 in maintaining and regaining stem cell pluripotency. Stem Cell Res. Ther. 1, 39. 10.1186/scrt39.

26. Liu, S., Cao, Y., Cui, K., Tang, Q., and Zhao, K. (2022). Hi-TrAC reveals division of labor of transcription factors in organizing chromatin loops. Nat. Commun. 13, 6679. 10.1038/s41467-022-34276-8.

27. Hua, P., Badat, M., Hanssen, L.L.P., Hentges, L.D., Crump, N., Downes, D.J., Jeziorska, D.M., Oudelaar, A.M., Schwessinger, R., Taylor, S., et al. (2021). Defining genome architecture at base-pair resolution. Nature 595, 125–129. 10.1038/s41586-021-03639-4.

28. Taylor, T., Sikorska, N., Shchuka, V.M., Chahar, S., Ji, C., Macpherson, N.N., Moorthy, S.D., de Kort, M.A.C., Mullany, S., Khader, N., et al. (2022). Transcriptional regulation and chromatin architecture maintenance are decoupled functions at the Sox2 locus. Genes Dev. 36, 699–717. 10.1101/gad.349489.122.

29. Sabari, B.R., Dall’Agnese, A., Boija, A., Klein, I.A., Coffey, E.L., Shrinivas, K., Abraham, B.J., Hannett, N.M., Zamudio, A.V., Manteiga, J.C., et al. (2018). Coactivator condensation at super-enhancers links phase separation and gene control. Science 361, eaar3958. 10.1126/science.aar3958.

30. Du, M., Stitzinger, S.H., Spille, J.-H., Cho, W.-K., Lee, C., Hijaz, M., Quintana, A., and Cissé, I.I. (2024). Direct observation of a condensate effect on super-enhancer controlled gene bursting. Cell 187, 331–344.e17. 10.1016/j.cell.2023.12.005.

31. Hnisz, D., Abraham, B.J., Lee, T.I., Lau, A., Saint-André, V., Sigova, A.A., Hoke, H.A., and Young, R.A. (2013). Super-Enhancers in the Control of Cell Identity and Disease. Cell 155, 934–947. 10.1016/j.cell.2013.09.053.

32. Cho, W.-K., Spille, J.-H., Hecht, M., Lee, C., Li, C., Grube, V., and Cisse, I.I. (2018). Mediator and RNA polymerase II clusters associate in transcription-dependent condensates. Science 361, 412–415. 10.1126/science.aar4199.

33. Mitchelmore, J., Grinberg, N.F., Wallace, C., and Spivakov, M. (2020). Functional effects of variation in transcription factor binding highlight long-range gene regulation by epromoters. Nucleic Acids Res. 48, 2866–2879. 10.1093/nar/gkaa123.

34. Reddy, T.E., Gertz, J., Pauli, F., Kucera, K.S., Varley, K.E., Newberry, K.M., Marinov, G.K., Mortazavi, A., Williams, B.A., Song, L., et al. (2012). Effects of sequence variation on differential allelic transcription factor occupancy and gene expression. Genome Res. 22, 860–869. 10.1101/gr.131201.111.

35. Gu, Z., Eils, R., and Schlesner, M. (2016). Complex heatmaps reveal patterns and correlations in multidimensional genomic data. Bioinformatics 32, 2847–2849. 10.1093/bioinformatics/btw313.

36. Palacio, M., and Taatjes, D.J. (2021). Merging established mechanisms with new insights: Condensates, hubs, and the regulation of RNA polymerase II transcription. J. Mol. Biol. 434, 167216. 10.1016/j.jmb.2021.167216.

37. Boehning, M., Dugast-Darzacq, C., Rankovic, M., Hansen, A.S., Yu, T., Marie-Nelly, H., McSwiggen, D.T., Kokic, G., Dailey, G.M., Cramer, P., et al. (2018). RNA polymerase II clustering through carboxy-terminal domain phase separation. Nat. Struct. Mol. Biol. 25, 833–840. 10.1038/s41594-018-0112-y.

38. Lim, B., and Levine, M.S. (2021). Enhancer-promoter communication: hubs or loops? Curr. Opin. Genet. Dev. 67, 5–9. 10.1016/j.gde.2020.10.001.

39. Ibrahim, A.Y., Khaodeuanepheng, N.P., Amarasekara, D.L., Correia, J.J., Lewis, K.A., Fitzkee, N.C., Hough, L.E., and Whitten, S.T. (2023). Intrinsically disordered regions that drive phase separation form a robustly distinct protein class. J. Biol. Chem. 299, 102801. 10.1016/j.jbc.2022.102801.

40. Bjarnason, S., McIvor, J.A.P., Prestel, A., Demény, K.S., Bullerjahn, J.T., Kragelund, B.B., Mercadante, D., and Heidarsson, P.O. (2024). DNA binding redistributes activation domain ensemble and accessibility in pioneer factor Sox2. Nat. Commun. 15, 1445. 10.1038/s41467-024-45847-2.

41. Sharma, R., Choi, K.-J., Quan, M.D., Sharma, S., Sankaran, B., Park, H., LaGrone, A., Kim, J.J., MacKenzie, K.R., Ferreon, A.C.M., et al. (2021). Liquid condensation of reprogramming factor KLF4 with DNA provides a mechanism for chromatin organization. Nat. Commun. 12, 5579. 10.1038/s41467-021-25761-7.

42. Chen, J., Zhang, Z., Li, L., Chen, B.-C., Revyakin, A., Hajj, B., Legant, W., Dahan, M., Lionnet, T., Betzig, E., et al. (2014). Single-molecule dynamics of enhanceosome assembly in embryonic stem cells. Cell 156, 1274–1285. 10.1016/j.cell.2014.01.062.

43. Lizio, M., Harshbarger, J., Shimoji, H., Severin, J., Kasukawa, T., Sahin, S., Abugessaisa, I., Fukuda, S., Hori, F., Ishikawa-Kato, S., et al. (2015). Gateways to the FANTOM5 promoter level mammalian expression atlas. Genome Biol. 16, 22. 10.1186/s13059-014-0560-6.

44. Cusanovich, D.A., Pavlovic, B., Pritchard, J.K., and Gilad, Y. (2014). The functional consequences of variation in transcription factor binding. PLoS Genet. 10, e1004226. 10.1371/journal.pgen.1004226.

45. Spivakov, M. (2014). Spurious transcription factor binding: non-functional or genetically redundant? BioEssays News Rev. Mol. Cell. Dev. Biol. 36, 798–806. 10.1002/bies.201400036.

46. Patel, Z.M., and Hughes, T.R. (2021). Global properties of regulatory sequences are predicted by transcription factor recognition mechanisms. Genome Biol. 22, 285. 10.1186/s13059-021-02503-y.

47. King, D.M., Hong, C.K.Y., Shepherdson, J.L., Granas, D.M., Maricque, B.B., and Cohen, B.A. (2020). Synthetic and genomic regulatory elements reveal aspects of cis-regulatory grammar in mouse embryonic stem cells. eLife 9, e41279. 10.7554/eLife.41279.

48. Hamley, J.C., Li, H., Denny, N., Downes, D., and Davies, J.O.J. (2023). Determining chromatin architecture with Micro Capture-C. Nat. Protoc. 18, 1687–1711. 10.1038/s41596-023-00817-8.

49. McLean, C.Y., Bristor, D., Hiller, M., Clarke, S.L., Schaar, B.T., Lowe, C.B., Wenger, A.M., and Bejerano, G. (2010). GREAT improves functional interpretation of cis-regulatory regions. Nat. Biotechnol. 28, 495–501. 10.1038/nbt.1630.

