## Supplemental Figues and Tables for "Enhancers mediate euchromatin hopping at chromatin contact points"

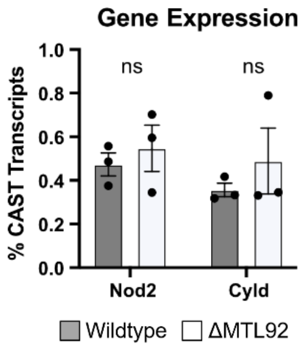


**Extended Data Figure 1 | Gene Expression of active genes within *Sall1* TAD after MTL92 Deletion**. *Nod2* and *Cyld* gene expression displayed as % CAST transcripts, calculated by (CAST reads / 129 + CAST reads) in MTL92 deleted cells. MTL92 deletion was done on the CAST allele only. Error bars represent SEM. No significant differences (ns) were observed (P>0.05).

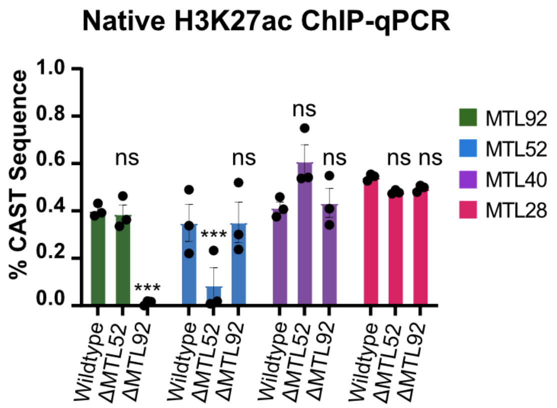


**Extended Data Figure 2 | Native H3K27ac enrichment after inactive MTLs are deleted.** Enrichment displayed as % CAST sequence, calculated by (CAST reads / 129 + CAST reads). All deletions were done on the CAST allele only. Error bars represent SEM. No significant differences = (ns), *= (P>0.05), ***= (P>0.01).

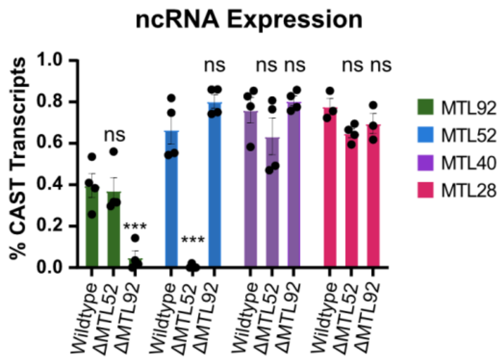


**Extended Data Figure 3 | ncRNA enrichment after inactive MTLs are deleted.** Enrichment displayed as % CAST sequence, calculated by (CAST reads / 129 reads + CAST reads). All deletions were done on the CAST allele only. Error bars represent SEM. No significant differences = (ns), *= (P>0.05), ***= (P>0.01).

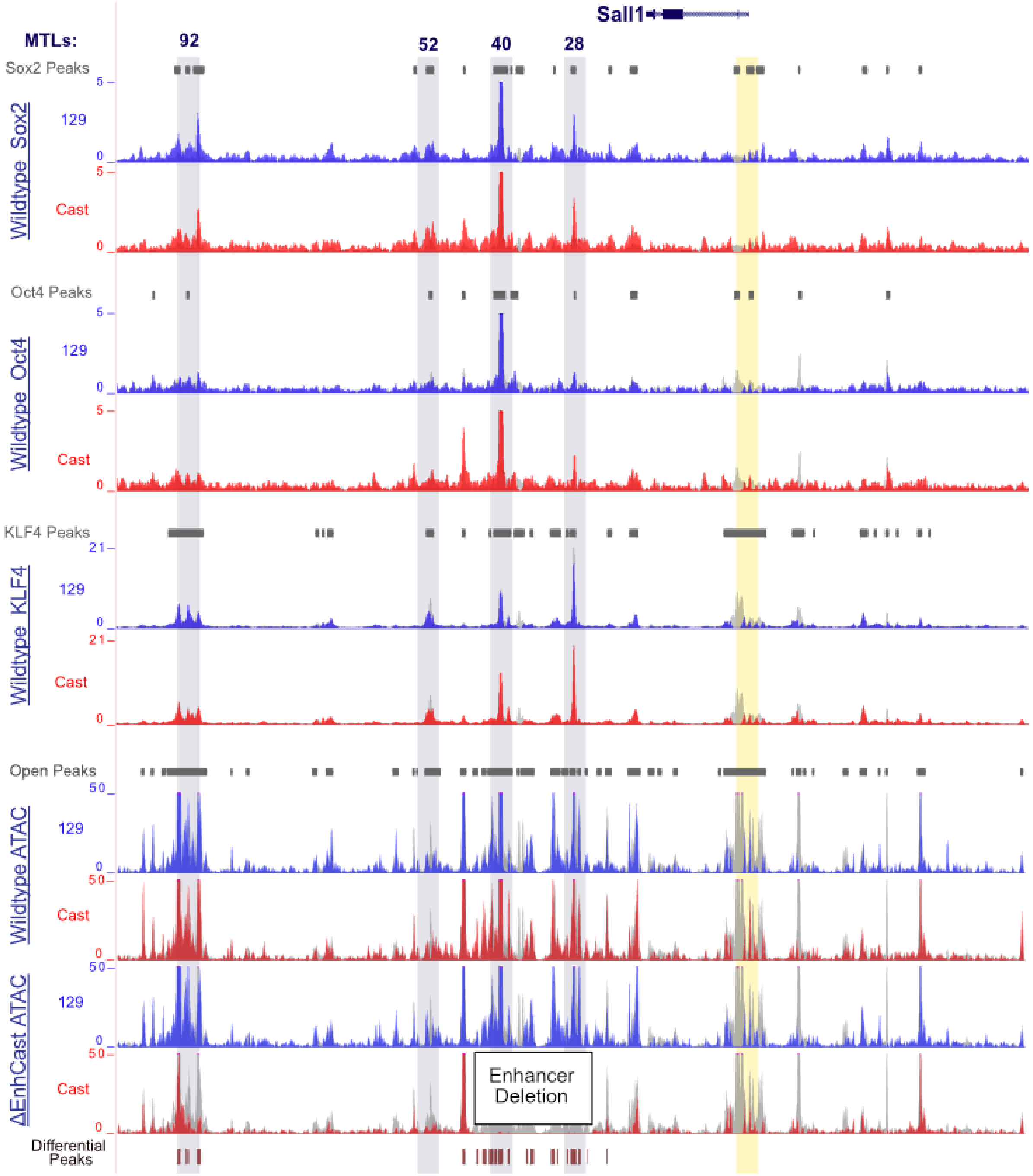


**Extended Data Figure 4 | Allele Specific OCT4, SOX2, and KLF4 binding tracks with open chromatin.** 129 reads are shown in blue, CAST in red, and total reads behind in grey. Each MTL is highlighted in grey with the Sall1 promoter in yellow. Peaks shown are calculated from all reads. All TF data is from wildtype cells and open chromatin data is from wildtype and ΔENHCAST clones, with the deletion coordinates represented in black box.

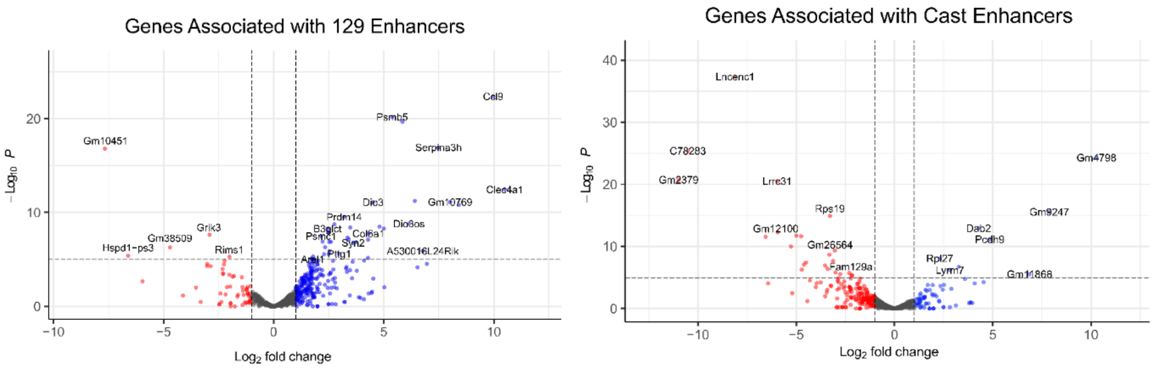


**Extended Data Figure 5 | Expression ratio of genes associated with allele specific enhancers.** Each allele specific enhancer was associated with an expressed gene in the same TAD and the expression ratio of 129 vs CAST was determined and visualized with EnhancedVolcano. The X-axis represents the fold change in RNA-seq reads between 129 and **CAST** while the Y-axis indicates statistical confidence, as shown by -log10(P-Value). Red dots represent genes with a significant differential expression favoring the CAST allele, and blue dots are differentially expressed genes favoring the 129 allele.

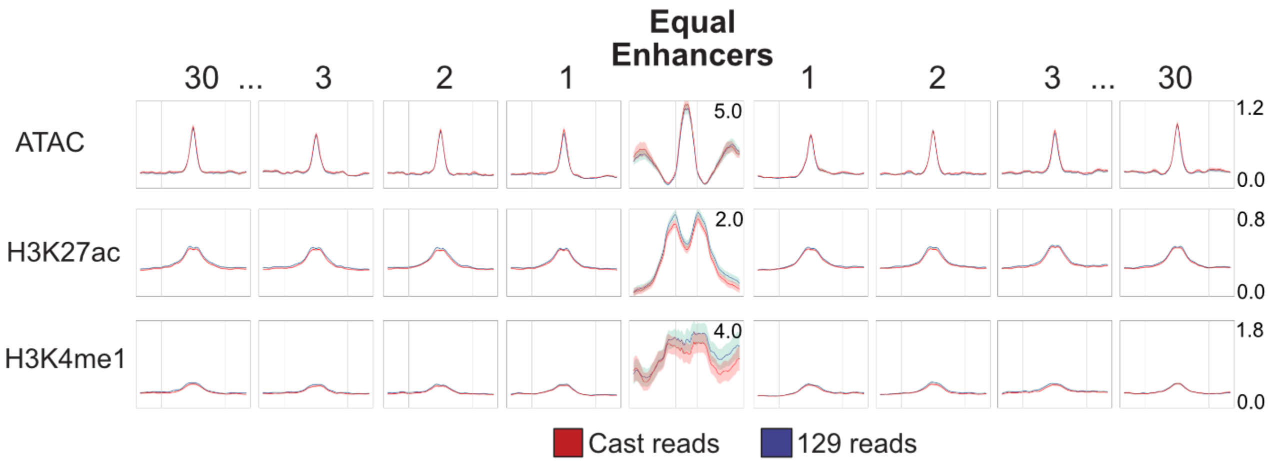


**Extended Data Figure 6 | Average profile plots of allele specific enhancers with high or low RNAPII signal.** NGS average profile plots of 129 and CAST reads for ATAC, H3K27ac, and H3K4me1 at ATAC peaks upstream and downstream enhancers with even levels of chromatin features on each allele. Average profile plots include a 4kb region centered on ATAC peaks and 1kb flank on either side. Peaks shown are the 1st , 2nd , 3rd, and 30th ATAC peak upstream and downstream from the enhancers.

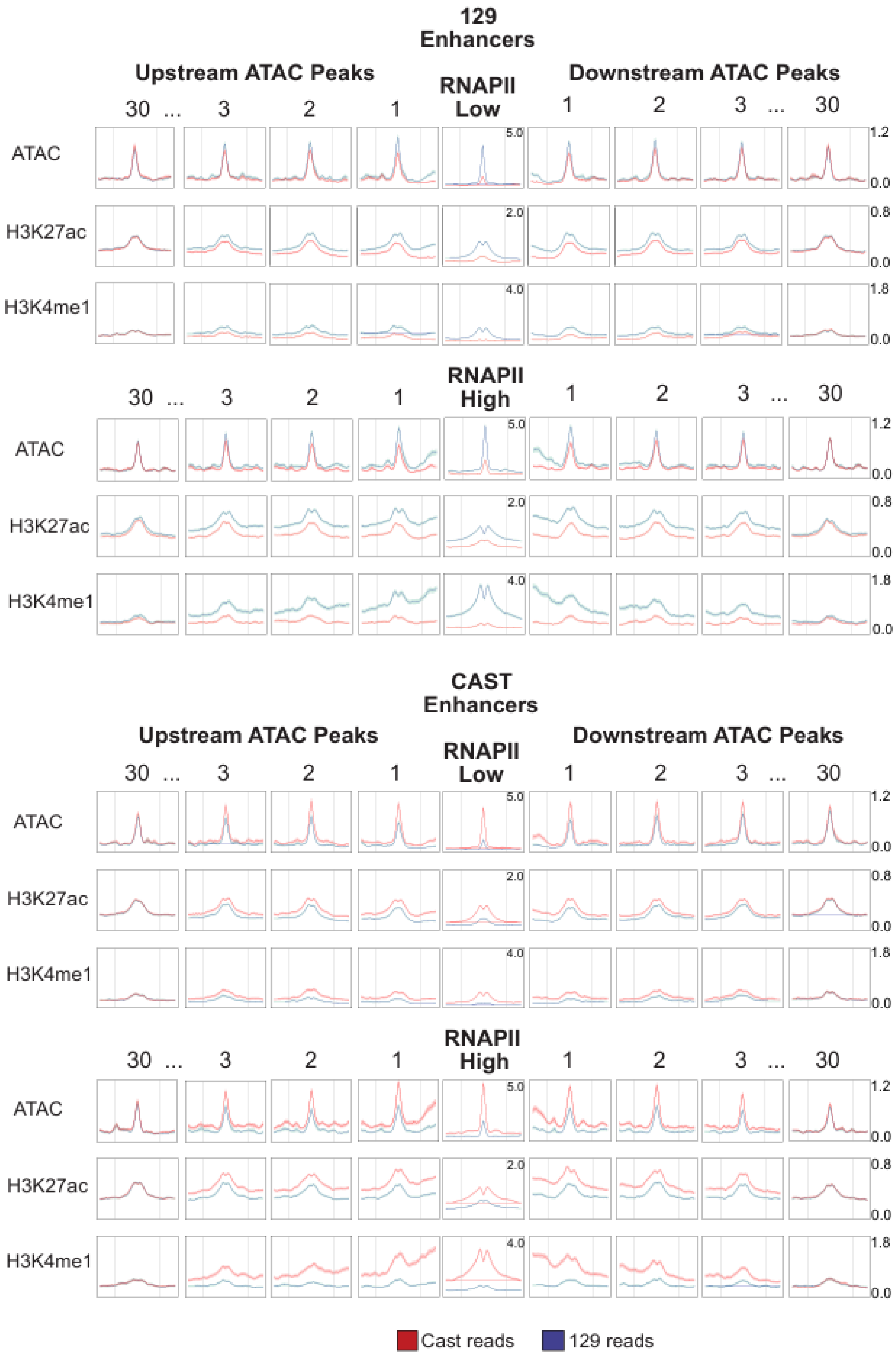


**Extended Data Figure 7 | Average profile plots of allele specific enhancers with high or low RNAPII signal.** NGS average profile plots of 129 and CAST reads for ATAC, H3K27ac, and H3K4me1 at ATAC peaks upstream and downstream of all 129 specific enhancers on top and CAST specific enhancers on the bottom. Enhancers were split up into those with more than 500 reads per million of RNAPII and those with less. Average profile plots include a 4kb region centered on ATAC peaks and 1kb flank on either side. Peaks shown are the 1st , 2nd , 3rd, and 30th ATAC peak upstream and downstream from the enhancers.

**Extended Data Table 1 | All primers and probes used in experiments**

| | Forward | Reverse |
|---|---|---|
| **gRNA** | | |
| MTL92 | ACGTGTGGCTGTATTAATAG | GTGGAGGACATTAGCCTCcG |
| MTL52 | TAAGCTGTGATGGCCTTATTGC | GAACCATTCTCATCTGACTCTGC |
| MTL40 | GAGTCTCTTCAGGACAACACCAT | GTCATCTAACAGCAAGCGAATCC |
| MTL28 | GGCACCTCTAGAAAGTAAGACCTG | AGCTACACAAAGGTGGGTATCATT |
| MTL28-40 | GAGTCTCTTCAGGACAACACCAT | AGCTACACAAAGGTGGGTATCATT |
| SCR | TAGCATACGTCACGCCGGAA | ACTGTTCTCGAACACTCTGT |
| **Gene Expression Primers** | | |
| Sall1_129 | CGTGGCCTTCTTGTCAATg | CAACAGTACTCTGAACTCCCCAgT |
| Sall1_CAST | CCGTGGCCTTCTTGTCAATa | CAACAGTACTCTGAACTCCCCAaT |
| cyld 129 | CTGTGTAGGTGTGAAGGAGTGGt | AGTGCACGCCACTGAACAT |
| cyld CAST | CAGGCCATGCCTGATACAT | GCAGATCCACACAGAGTCAGTAg |
| nod2 129 | TAACTGAACGCCTGTTGCTG | GTGGAGGTATCAGAGTGGGATGa |
| nod2 CAST | AACTGAACGCCTGTTGCTGT | GGAGGTATCAGAGTGGGATGg |
| **eRNA Primers** | | |
| 52_129 | GAGGTTCAAAGGGTCACAACC | AATTTGCCTCGTATTAATTTTGCT |
| 52_CAST | GAGGTTCAAAGGGTCACAACt | ATGCTAAGCAATGGCAGAGG |
| 92_129 | TTAAGCCTGGTAGCTTGAAGAg | TTCCCGAATGACCCTAGTTG |
| 92_CAST | TTAAGCCTGGTAGCTTGAAGAa | TTCCCGAATGACCCTAGTTG |
| 28_129 | TCAAGGTATGGCCTCTGAaa | CTCTCTCGGGTGGCTGCTAT |
| 28_CAST | TCAAGGTATGGCCTCTGAag | CTCTCTCGGGTGGCTGCTAT |
| 40_129 | TCCTCTTAACCCCAGGGACt | GCAAGTCAAGTTGTACCCACA |
| 40_CAST | TCCTCTTAACCCCAGGGACc | GCAAGTCAAGTTGTACCCACA |
| **TF ChIP Primers** | | |
| 92 129 | CCTAAGGCACCATCACAGGTc | GCCTTTGTGTGTTTGGCTTT |
| 92 CAST | AGGCTCTTTGGAAGTCTTGGAt | AAGCAAGCCACACCTTTGTC |
| 52 129 | AAGGCTCCAGAAAGGCAAGT | CACGCAGAGGATGGGGTAa |
| 52CAST | AAGGCTCCAGAAAGGCAAGT | CACGCAGAGGATGGGGTAg |
| **Sox2 insert Primers** | | |
| Inside | CCACACTACGCGGACTTTTT | TGGTGCCCTATAGCTCGTTT |
| Upstream | CAATATGCCTGCTTtCTTCAc | TCCACCGTTTAGGTAATGACG |
| Downstream | CGGcTAGACGTGGATGAACt | CAAACAGTGCTCaCGCAGAC |
| Positive Control | CAGGCAGGAAAGAGAAGGTG | CCCAGATTCCAGGCTTTGTA |
| Negative Control | cGTGTGTTGTTTCCTCTCCt | CTGCCTTCCTACAGCTCCAC |
| **MicroCapture-C Probes** | | |
| Promoter_1 | CCCCGGGAACGTGATCAGACCGGAACACTTTTAAATCTAGCACTCTCCC<br>CTACCCCGCAAATCGGAGCGCTTGAAAACACACCAACGTGCACCCCTCA<br>GGGTGCACCGAAGCTTCCAGGC | |
| Promoter_2 | CCAGAGCTGGGAGCCAGCCTCTCTTACCCGCTCACGGGACCCAGGGAT<br>CCGGGCCCTGTACTCCCTCTTCCGGGTTTGGGCGCCTGCGAACCTGAGG<br>CTTGGTCCTCGGCTTAGTTTAGA | |

All lower case letters indicate a SNP between 129 and CAST alleles.

**Extended Data Table 2 | Differential H3K27ac peaks after enhancer deletion**

| Chr | Start | End | P-value | FDR | Log2 Fold Change | Shrunk Log2 Fold Change |
|---|---|---|---|---|---|---|
| 8 | 89016397 | 89022616 | 0 | 0 | 4.986973 | 4.215974 |
| 8 | 88988247 | 89016206 | 0 | 0 | 4.772696 | 4.585989 |
| 8 | 89036681 | 89038411 | 8.39E-07 | 3.03E-05 | 3.159214 | 0.667034 |
| 8 | 88945463 | 88955273 | 0 | 0 | 2.87683 | 2.749717 |
| 8 | 89023301 | 89028940 | 0 | 0 | 2.806413 | 2.436065 |
| 8 | 89039074 | 89048113 | 0 | 0 | 2.783881 | 2.447407 |
| 9 | 52801461 | 52805761 | 0 | 0 | 2.566994 | 2.1848 |
| 7 | 1.42E+08 | 1.42E+08 | 2.37E-06 | 7.66E-05 | 2.554432 | 0.782479 |
| 8 | 89062981 | 89065696 | 1.11E-35 | 1.03E-32 | 2.435818 | 1.842384 |
| 8 | 88944517 | 88944872 | 2.05E-04 | 0.00361 | 2.383064 | 0.55915 |
| 8 | 88985420 | 88986510 | 7.29E-08 | 3.39E-06 | 2.345841 | 0.921386 |
| 2 | 45695897 | 45702903 | 0 | 2.24E-44 | 2.329186 | 1.907046 |
| 6 | 43442438 | 43443019 | 9.64E-05 | 0.0019 | 2.295972 | 0.611084 |
| 2 | 1.62E+08 | 1.62E+08 | 3.34E-29 | 1.86E-26 | 2.29563 | 1.695324 |
| 4 | 52585871 | 52592030 | 0 | 1.40E-45 | 2.279724 | 1.888049 |
| 10 | 78568914 | 78569384 | 3.81E-06 | 1.17E-04 | 2.246729 | 0.7715 |
| 8 | 88972293 | 88976294 | 0 | 0 | 2.195675 | 1.859624 |
| 11 | 35077669 | 35083891 | 0 | 0 | 2.12552 | 1.871528 |
| 17 | 79397205 | 79398570 | 1.70E-24 | 6.41E-22 | 2.015976 | 1.510756 |
| 9 | 62649562 | 62650117 | 3.27E-07 | 1.28E-05 | 2.01434 | 0.879489 |

H3K27ac Native ChIP reads from wildtype CAST allele and enhancer deleted CAST allele were compared and differential peaks with cut off of $P<0.05$ and Log 2 fold change >2 was calculated with DeSeq to determine effects of enhancer deletion.

**Extended Data Table 3 | Transcription factor motif scores within MTL sequences**

MTL28

| Model ID | Model name | Relative score | Start | End | Strand | predicted site sequence |
|---|---|---|---|---|---|---|
| MA0039.2 | Klf4 | 0.998424651771266 | 1065 | 1074 | 1 | TGGGTGTGGC |
| MA0143.3 | Sox2 | 0.971704544250396 | 1142 | 1149 | 1 | CCATTGTG |
| MA0792.1 | POU5F1B | 0.844229229456798 | 1415 | 1423 | 1 | AATGCTAAC |

MTL40

| Model ID | Model name | Relative score | Start | End | Strand | predicted site sequence |
|---|---|---|---|---|---|---|
| MA0039.2 | Klf4 | 0.959585936546979 | 10754 | 10763 | 1 | GGGGTGGGGT |
| MA0143.3 | Sox2 | 0.980290280623661 | 293 | 300 | 1 | CCATTGTC |
| MA0792.1 | POU5F1B | 1.00000793573648 | 4630 | 4638 | 1 | TATGCAAAT |

MTL52

| Model ID | Model name | Relative score | Start | End | Strand | predicted site sequence |
|---|---|---|---|---|---|---|
| MA0039.2 | Klf4 | 0.959585936546979 | 1382 | 1391 | 1 | GGGGTGGGGT |
| MA0143.3 | Sox2 | 0.887816223363677 | 717 | 724 | -1 | CCTTTGTA |
| MA0792.1 | POU5F1B | 0.918225600810559 | 1255 | 1263 | 1 | AATGTAAAT |

MTL92

| Model ID | Model name | Relative score | Start | End | Strand | predicted site sequence |
|---|---|---|---|---|---|---|
| MA0039.2 | Klf4 | 0.993956330640913 | 3260 | 3269 | -1 | AGGGTGTGGC |
| MA0143.3 | Sox2 | 0.985026626928747 | 4256 | 4263 | 1 | CCTTTGTC |
| MA0792.1 | POU5F1B | 0.926467231462008 | 5454 | 5462 | -1 | CATGTAAAT |

JASPAR 2016 was used to find Oct4 (POU5F1B), Sox2, and KLF4 motifs and their score representing % match with motif, and location of match within the sequence of MTL28, 40, 52, and 92.

**Extended Data Table 4 | Average profile plot distance and adjusted p-value for 129 and CAST ATAC, H3K27ac, and H3K4me1 read counts.**

**ATAC peaks upstream of CAST enhancers**

| Peaks away | ATAC Q-Value | K27ac Q-Value | K4me1 Q-Value | Distance |
|---|---|---|---|---|
| 1 | 7.29E-70 | 4.20E-71 | 2.60E-159 | 16323.994 |
| 2 | 2.62E-33 | 1.17E-47 | 3.74E-111 | 27675.428 |
| 3 | 1.93E-28 | 4.06E-36 | 9.02E-85 | 38636.086 |
| 4 | 1.45E-23 | 1.01E-27 | 5.98E-70 | 50568.195 |
| 5 | 2.82E-19 | 8.53E-22 | 4.52E-51 | 62320.278 |
| 6 | 1.25E-12 | 2.83E-17 | 4.28E-44 | 74397.009 |
| 7 | 3.35E-10 | 1.34E-14 | 1.27E-39 | 86936.667 |
| 8 | 1.46E-12 | 8.72E-17 | 1.26E-39 | 98167.83 |
| 9 | 1.24E-15 | 3.29E-15 | 1.47E-36 | 109672.33 |
| 10 | 5.67E-09 | 1.90E-13 | 2.66E-29 | 121046.83 |
| 11 | 0.0016748 | 8.53E-05 | 2.11E-16 | 135193.39 |
| 12 | 20.752478 | 4.76E-05 | 1.90E-13 | 147027.44 |
| 13 | 0.905247 | 8.47E-07 | 6.34E-14 | 160030.82 |
| 14 | 0.0163492 | 2.46E-07 | 6.76E-15 | 171222.18 |
| 15 | 0.3862533 | 3.96E-06 | 2.83E-13 | 182690.26 |
| 16 | 0.0005336 | 7.93E-07 | 2.95E-16 | 194871.19 |
| 17 | 0.0166129 | 1.22E-05 | 1.84E-11 | 206713.63 |
| 18 | 0.0755033 | 2.85E-05 | 4.79E-12 | 218746.63 |
| 19 | 0.2067298 | 0.0002065 | 3.26E-11 | 230698.55 |
| 20 | 0.5637353 | 0.0004615 | 4.34E-10 | 243056.95 |
| 21 | 48.546862 | 3.8379142 | 3.76E-06 | 254171.72 |
| 22 | 25.529548 | 0.1142901 | 2.78E-06 | 265939.72 |
| 23 | 0.53791 | 0.0115925 | 0.00012264 | 277844.74 |
| 24 | 669.74033 | 3.9177092 | 7.21E-05 | 289622.58 |
| 25 | 30.595958 | 2.3659839 | 7.03E-06 | 301384.96 |
| 26 | 0.776481 | 1.8719027 | 6.24E-10 | 312714.45 |
| 27 | 8.7544119 | 4.3195699 | 1.86E-07 | 324047.99 |
| 28 | 1069.7943 | 83.445246 | 0.00572619 | 336044.03 |
| 29 | 724.60757 | 41.867375 | 0.05052963 | 347100.49 |
| 30 | 797.83028 | 122.66342 | 0.00817245 | 359230.06 |
| 31 | 349.7821 | 207.95336 | 0.70646259 | 371862.22 |
| 32 | 191.68273 | 631.02102 | 0.39909059 | 382919.22 |
| 33 | 954.86822 | 814.5704 | 1.35238293 | 393739.26 |
| 34 | 47.941235 | 160.71653 | 0.34483074 | 404808.52 |
| 35 | 93.23638 | 364.22737 | 0.0256564 | 415860.09 |
| 36 | 1835.3064 | 360.2662 | 0.00194215 | 426401.6 |
| 37 | 275.61649 | 65.87084 | 0.02694144 | 436940.96 |
| 38 | 46.585352 | 197.36265 | 2.36143789 | 448051.91 |
| 39 | 73.234372 | 172.07233 | 275.444248 | 457958.53 |
| 40 | 0.2837822 | 1.14E+00 | 1.32E+01 | 469155.66 |

**ATAC peaks downstream of CAST enhancers**

| Peaks away | ATAC Q-Value | K27ac Q-Value | K4me1Q-Value | Distance |
|---|---|---|---|---|
| 1 | 1.92E-55 | 5.62E-66 | 2.53E-149 | 15582.17607 |
| 2 | 1.07E-33 | 1.62E-51 | 1.30E-102 | 26275.83014 |
| 3 | 2.06E-17 | 1.58E-35 | 3.20E-80 | 37359.79803 |
| 4 | 3.51E-19 | 8.26E-26 | 1.09E-68 | 48474.25117 |
| 5 | 1.71E-12 | 2.06E-17 | 2.56E-43 | 60285.35111 |
| 6 | 2.70E-11 | 2.90E-15 | 2.57E-37 | 71594.70016 |
| 7 | 6.44E-12 | 2.84E-12 | 1.31E-35 | 82708.78716 |
| 8 | 1.43E-05 | 3.12E-10 | 7.46E-27 | 94658.08389 |
| 9 | 1.55E-07 | 4.44E-10 | 3.20E-28 | 107051.9063 |
| 10 | 2.31E-05 | 7.31E-11 | 3.63E-29 | 118282.8446 |
| 11 | 9.3801308 | 2.90E-07 | 1.06E-25 | 132964.465 |
| 12 | 0.0803148 | 0.001044 | 3.54E-13 | 145816.5215 |
| 13 | 0.0231122 | 3.92E-06 | 1.40E-16 | 157675.0746 |
| 14 | 0.1920837 | 6.61E-07 | 6.08E-19 | 169705.6649 |
| 15 | 0.0179982 | 3.62E-07 | 1.39E-18 | 180619.3195 |
| 16 | 0.1345264 | 0.0066012 | 3.50E-12 | 192666.638 |
| 17 | 1.5424607 | 0.0728299 | 2.27E-08 | 203992.1129 |
| 18 | 0.4653973 | 0.0025357 | 6.26E-07 | 215141.9907 |
| 19 | 2.1421433 | 4.27E-05 | 2.85E-05 | 226981.9715 |
| 20 | 0.3391327 | 0.0011175 | 0.000040767 | 237967.6808 |
| 21 | 699.99653 | 0.4197075 | 0.00610755 | 249052.911 |
| 22 | 170.087 | 0.0761162 | 0.101791187 | 260914.9824 |
| 23 | 49.693549 | 0.5514916 | 0.000111583 | 272279.0264 |
| 24 | 636.50173 | 23.68996 | 0.001557091 | 283361.0367 |
| 25 | 14.806539 | 33.597768 | 0.000332314 | 294918.2007 |
| 26 | 1.42E+02 | 414.34613 | 0.001965701 | 305517.2043 |
| 27 | 13.878863 | 54.655713 | 0.002024122 | 316270.732 |
| 28 | 1.61E+02 | 81.621487 | 0.168441786 | 327492.208 |
| 29 | 27.980183 | 15.365174 | 0.064939052 | 339066.7377 |
| 30 | 1.10E+03 | 300.3625 | 0.1403985 | 350764.1309 |
| 31 | 961.69976 | 184.51286 | 1.811585572 | 365294.1141 |
| 32 | 1.05E+03 | 951.46921 | 153.8707068 | 377773.9616 |
| 33 | 944.16422 | 1294.4155 | 16.97572636 | 389369.2199 |
| 34 | 1.41E+03 | 837.40943 | 1.043296295 | 400957.1323 |
| 35 | 76.739924 | 10.44401 | 0.006720992 | 412389.7905 |
| 36 | 1.39E+02 | 14.655115 | 0.019708265 | 423848.1234 |
| 37 | 48.463433 | 7.0908849 | 7.127037198 | 435564.7697 |
| 38 | 2.96E-01 | 0.1323373 | 0.008941502 | 446967.6437 |
| 39 | 1.5770632 | 0.0235789 | 0.031436905 | 459078.9123 |
| 40 | 2.86E+02 | 2.21E-02 | 1.84E-01 | 469745.8377 |

**ATAC peaks upstream of 129 Enhancers**

| Peaks away | ATAC Q-Value | K27ac Q-Value | K4me1 Q-Value | Distance |
|---|---|---|---|---|
| 1 | 1.55E-80 | 9.10E-72 | 9.02E-155 | 17978.26752 |
| 2 | 1.86E-60 | 9.91E-58 | 1.74E-116 | 29441.59551 |
| 3 | 8.59E-44 | 1.04E-44 | 1.93E-101 | 41591.18513 |
| 4 | 1.76E-45 | 9.03E-35 | 6.33E-83 | 53279.13055 |
| 5 | 9.29E-37 | 1.37E-25 | 1.69E-70 | 65897.397 |
| 6 | 5.72E-30 | 3.92E-20 | 1.64E-54 | 78203.35045 |
| 7 | 1.90E-25 | 1.43E-20 | 2.35E-51 | 91115.37079 |
| 8 | 4.27E-20 | 4.54E-14 | 3.25E-33 | 102817.4312 |
| 9 | 8.70E-15 | 2.22E-10 | 3.25E-34 | 115336.061 |
| 10 | 5.58E-13 | 1.70E-14 | 1.61E-31 | 127405.9486 |
| 11 | 1.38E-08 | 1.04E-09 | 5.33E-26 | 142178.6977 |
| 12 | 1.68E-08 | 4.67E-10 | 1.35E-22 | 154570.0999 |
| 13 | 3.71E-15 | 2.80E-08 | 2.29E-22 | 167537.4697 |
| 14 | 9.92E-17 | 5.63E-07 | 7.02E-23 | 180334.0744 |
| 15 | 2.91E-10 | 0.037891 | 3.62E-20 | 192763.5531 |
| 16 | 6.81E-05 | 5.7250544 | 5.74E-14 | 204773.8183 |
| 17 | 0.0006508 | 0.0428065 | 2.33E-11 | 217502.9176 |
| 18 | 0.0007891 | 0.1296685 | 3.65E-12 | 229727.6217 |
| 19 | 0.0001026 | 3.5880117 | 3.08E-10 | 242120.4431 |
| 20 | 4.06E-05 | 0.6188302 | 6.75E-09 | 254267.3114 |
| 21 | 0.0002878 | 0.0027951 | 1.99E-09 | 265814.8443 |
| 22 | 1.95E-05 | 0.0173553 | 5.97E-09 | 277668.949 |
| 23 | 0.0079415 | 63.507197 | 5.32E-08 | 289175.2136 |
| 24 | 0.0006484 | 7.3113201 | 0.00026054 | 301351.8533 |
| 25 | 1.49E-07 | 0.6158344 | 0.00020066 | 313075.848 |
| 26 | 6.59E-06 | 0.0093464 | 2.94E-07 | 324403.6026 |
| 27 | 0.0672793 | 0.688559 | 6.12E-05 | 335423.6711 |
| 28 | 0.0306326 | 8.356503 | 1.04E-05 | 347294.8066 |
| 29 | 0.0411508 | 298.82371 | 0.00942777 | 359646.4166 |
| 30 | 0.4641221 | 262.17392 | 1.012275 | 371729.8135 |
| 31 | 10.429201 | 600.72916 | 39.1007508 | 384987.9048 |
| 32 | 10.573961 | 1093.7888 | 28.3837248 | 396834.7707 |
| 33 | 1.329299 | 911.73371 | 8.96201818 | 408269.2878 |
| 34 | 12.345367 | 1519.8879 | 40.5090407 | 419589.9941 |
| 35 | 0.0643987 | 1458.4028 | 3.44616354 | 431303.4787 |
| 36 | 2.4807044 | 1429.8944 | 312.243937 | 442787.1356 |
| 37 | 1.8609561 | 1283.9509 | 802.687857 | 454732.4734 |
| 38 | 32.988016 | 1470.633 | 1316.28307 | 465930.0165 |
| 39 | 0.0001916 | 172.6656 | 1.10298548 | 477276.9021 |
| 40 | 21.06307 | 8.13E+02 | 7.18E+01 | 488910.6431 |

**ATAC peaks downstream of 129 Enhancers**

| Peaks away | ATAC Q-Value | K27ac Q-Value | K4me1 Q-Value | Distance |
|---|---|---|---|---|
| 1 | 2.88E-85 | 4.07E-74 | 3.80E-170 | 18456.61301 |
| 2 | 3.76E-61 | 5.04E-51 | 1.81E-108 | 30910.61407 |
| 3 | 4.83E-49 | 1.53E-38 | 3.70E-84 | 43977.84382 |
| 4 | 5.05E-43 | 4.34E-30 | 2.27E-64 | 56745.11887 |
| 5 | 2.41E-37 | 8.58E-27 | 4.39E-65 | 69775.40192 |
| 6 | 1.35E-33 | 4.79E-23 | 1.52E-59 | 82097.16738 |
| 7 | 1.90E-14 | 1.01E-15 | 1.53E-30 | 94562.39126 |
| 8 | 1.24E-15 | 5.12E-12 | 8.48E-24 | 106271.4488 |
| 9 | 1.51E-11 | 3.39E-06 | 6.22E-20 | 118540.7292 |
| 10 | 6.53E-06 | 0.0294635 | 1.32E-13 | 130875.6679 |
| 11 | 1.43E-10 | 0.0001311 | 7.07E-26 | 145344.2839 |
| 12 | 1.84E-14 | 6.48E-07 | 4.69E-24 | 158723.3222 |
| 13 | 5.07E-16 | 1.97E-07 | 7.46E-27 | 170894.2674 |
| 14 | 2.52E-11 | 0.0002766 | 9.08E-24 | 183671.4104 |
| 15 | 5.25E-12 | 0.0057456 | 7.79E-18 | 195869.4375 |
| 16 | 2.61E-07 | 0.0007562 | 3.25E-16 | 209089.7018 |
| 17 | 0.0023608 | 0.0114904 | 4.79E-19 | 221699.8761 |
| 18 | 0.0006215 | 0.000605 | 2.08E-17 | 234608.16 |
| 19 | 3.23E-07 | 0.0611569 | 1.69E-15 | 247017.1388 |
| 20 | 6.70E-06 | 0.000454 | 1.34E-12 | 260725.4713 |
| 21 | 7.10E-08 | 0.0004531 | 8.04E-11 | 273619.1334 |
| 22 | 1.22E-08 | 0.0021579 | 2.40E-06 | 286196.9655 |
| 23 | 6.75E-12 | 0.283634 | 7.17E-11 | 298797.3443 |
| 24 | 3.35E-05 | 18.881368 | 0.00362287 | 310958.4283 |
| 25 | 0.0007136 | 21.637712 | 0.01381431 | 323763.5213 |
| 26 | 4.84E-08 | 1.8276221 | 0.00394034 | 335980.4107 |
| 27 | 4.21E-07 | 4.6420458 | 0.00299372 | 348395.4516 |
| 28 | 9.8170596 | 253.56465 | 0.40623363 | 361027.0399 |
| 29 | 1.0291556 | 594.46341 | 0.03334909 | 372740.6105 |
| 30 | 0.1389566 | 344.21224 | 2.21136075 | 384521.4182 |
| 31 | 0.560714 | 1219.5797 | 6.94393677 | 398639.3684 |
| 32 | 0.0014559 | 8.021534 | 0.03644045 | 409988.7087 |
| 33 | 0.0102726 | 40.164476 | 1.31661996 | 421712.9389 |
| 34 | 0.1891663 | 45.086555 | 0.13156863 | 433973.8357 |
| 35 | 0.2962547 | 1632.9826 | 7.29E-06 | 445870.9006 |
| 36 | 0.1180543 | 671.04219 | 0.00081885 | 458039.3636 |
| 37 | 0.0245821 | 403.90129 | 4.52E-05 | 469709.2828 |
| 38 | 1.38E-05 | 38.048781 | 9.63E-06 | 481780.5056 |
| 39 | 5.12E-11 | 32.156925 | 0.00180854 | 493160.0893 |
| 40 | 0.3500145 | 284.30875 | 12.2400981 | 504974.1893 |

For each ATAC peak upstream and downstream allele specific enhancers we calculated the adjusted P-Value (Q-Value) of 129 and CAST read counts for ATAC, H3K27ac, and H3K4me1 data. The last column is the peaks distance from the allele specific enhancer.

**Extended Data Table 5 | Average profile plot distance and adjusted p-value for 129 and CAST ATAC, H3K27ac, and H3K4me1 read counts at equal enhancers**

**ATAC peaks downstream of enhancers equal on each allele**

| Peaks away | ATAC Q-Value | K27ac Q-Value | K4me1 Q-value | Distance |
|---|---|---|---|---|
| 1 | 1745.64 | 252.9195 | 68.4025 | 8663.52 |
| 2 | 815.4311 | 26.76729 | 12.35568 | 17449.34 |
| 3 | 677.403 | 34.55033 | 6.538218 | 27150.31 |
| 4 | 1802.776 | 217.5987 | 2.189734 | 36184.62 |
| 5 | 173.2992 | 894.5302 | 687.3663 | 45665.56 |
| 6 | 84.32329 | 1468.072 | 1657.433 | 54131.16 |
| 7 | 796.9907 | 1565.499 | 1675.576 | 63084.62 |
| 8 | 1441.017 | 1839.306 | 1607.255 | 72656.41 |
| 9 | 902.3887 | 1438.137 | 227.8453 | 81417.48 |
| 10 | 180.6263 | 270.3192 | 66.07752 | 90388.26 |
| 11 | 1342.514 | 838.3662 | 144.7312 | 99653.37 |
| 12 | 692.6139 | 408.0096 | 40.88732 | 108634.4 |
| 13 | 323.3307 | 230.86 | 11.05352 | 117756 |
| 14 | 771.9646 | 1746.669 | 563.4419 | 126855 |
| 15 | 92.70906 | 1693.785 | 1472.861 | 136400.9 |
| 16 | 1139.532 | 754.2059 | 45.09947 | 145127.8 |
| 17 | 848.353 | 48.81309 | 34.6622 | 154597.8 |
| 18 | 681.6443 | 14.86534 | 128.3245 | 163678.1 |
| 19 | 888.7999 | 158.0741 | 347.2414 | 173048.8 |
| 20 | 1124.278 | 102.9903 | 81.63819 | 182141.4 |
| 21 | 1204.092 | 84.71237 | 125.3557 | 191929.1 |
| 22 | 1758.784 | 428.8541 | 373.7213 | 201324.9 |
| 23 | 1356.384 | 178.7937 | 600.8996 | 210270.1 |
| 24 | 1480.92 | 719.4954 | 848.1104 | 219400.9 |
| 25 | 565.1722 | 1698.396 | 973.1702 | 228741.6 |
| 26 | 630.1232 | 543.0766 | 452.2691 | 238142.1 |
| 27 | 532.0457 | 334.6954 | 899.0495 | 248120.8 |
| 28 | 1929.2 | 1253.971 | 666.4086 | 257643 |
| 29 | 834.0448 | 1715.613 | 426.2997 | 266696 |
| 30 | 1129.322 | 1310.054 | 1175.531 | 276174.5 |
| 31 | 1651.723 | 691.4711 | 1372.253 | 285916.8 |
| 32 | 108.8691 | 825.3112 | 655.9973 | 294942.9 |
| 33 | 2.747409 | 273.3399 | 934.6621 | 304170.4 |
| 34 | 377.7288 | 1221.428 | 1646.503 | 312823.6 |
| 35 | 929.8367 | 709.3808 | 209.576 | 322599.3 |
| 36 | 1713.536 | 215.4802 | 89.4231 | 332425.9 |
| 37 | 568.081 | 391.5828 | 464.5637 | 341961 |
| 38 | 1883.89 | 1074.409 | 406.3916 | 351931.7 |
| 39 | 153.7963 | 322.1724 | 1382.823 | 361422.3 |
| 40 | 24.60422 | 464.8613 | 1394.776 | 370850.4 |

**ATAC peaks upstream of enhancers equal on each allele**

| Peaks away | ATAC Q-Value | K27ac Q-Value | K4me1 Q-value | Distance |
|---|---|---|---|---|
| 1 | 1325.461 | 1459.221 | 333.0406 | 9320.38 |
| 2 | 1398.111 | 125.1284 | 39.27346 | 18827.27 |
| 3 | 1252.932 | 32.9668 | 2.125214 | 28800.53 |
| 4 | 702.1938 | 92.44414 | 1.644947 | 38011.7 |
| 5 | 294.1171 | 158.0984 | 21.65427 | 46939.04 |
| 6 | 1364.26 | 344.397 | 727.5759 | 56178.26 |
| 7 | 1796.735 | 539.6105 | 211.3117 | 65767.14 |
| 8 | 193.6537 | 1939.377 | 229.2343 | 74835.75 |
| 9 | 453.5723 | 1829.898 | 1628.467 | 84070.57 |
| 10 | 256.916 | 688.3647 | 1210.805 | 93438.49 |
| 11 | 375.3839 | 1440.808 | 1935.033 | 102436 |
| 12 | 1656.658 | 1874.246 | 1947.257 | 111951.4 |
| 13 | 429.1529 | 201.8599 | 123.1344 | 121878.9 |
| 14 | 455.1863 | 174.4357 | 89.06621 | 130921.9 |
| 15 | 1665.952 | 191.723 | 95.60569 | 139690.6 |
| 16 | 464.782 | 1899.722 | 116.2367 | 149056.6 |
| 17 | 136.8184 | 1561.564 | 62.4692 | 158582.7 |
| 18 | 997.7494 | 396.3757 | 4.434431 | 167841 |
| 19 | 793.8929 | 148.5327 | 2.466185 | 177578.8 |
| 20 | 1640.117 | 705.8971 | 47.18485 | 187070.2 |
| 21 | 1810.227 | 176.8517 | 35.79954 | 196246.7 |
| 22 | 1316.051 | 20.53844 | 39.92301 | 205643.4 |
| 23 | 1617.124 | 34.48278 | 1.121573 | 215184.3 |
| 24 | 1557.473 | 197.4005 | 56.75201 | 224367.4 |
| 25 | 1403.574 | 427.8231 | 19.18174 | 233577.3 |
| 26 | 163.8752 | 10.89848 | 45.46459 | 242811.9 |
| 27 | 1182.11 | 245.4086 | 105.3971 | 251999.8 |
| 28 | 489.7589 | 738.5192 | 256.8857 | 262080.9 |
| 29 | 679.2239 | 94.31391 | 14.92965 | 271729.8 |
| 30 | 1087.078 | 152.6355 | 32.47879 | 280972.3 |
| 31 | 1928.688 | 48.89058 | 49.60153 | 290727.8 |
| 32 | 1673.736 | 144.0757 | 169.1613 | 300117 |
| 33 | 711.876 | 1799.569 | 173.4878 | 309561.1 |
| 34 | 255.713 | 1339.21 | 640.1235 | 319348.2 |
| 35 | 635.7113 | 1885.361 | 98.74055 | 329044.2 |
| 36 | 1435.277 | 946.2843 | 342.0996 | 338588.6 |
| 37 | 1633.812 | 1597.532 | 580.14 | 347981.7 |
| 38 | 1386.866 | 469.7836 | 365.459 | 357731.7 |
| 39 | 1842.916 | 615.2728 | 701.5674 | 366798.3 |
| 40 | 1659.634 | 1939.371 | 938.4584 | 376246.7 |

For each ATAC peak upstream and downstream of enhancers with equal ATAC signal on each allele, we calculated the adjusted P-Value (Q-Value) of 129 and CAST read counts for ATAC, H3K27ac, and H3K4me1 data. The last column is the peaks distance from the allele specific enhancer.

**Extended Data Table 6 | Average profile plot distance and adjusted p-value for 129 and CAST ATAC, H3K27ac, and H3K4me1 read counts at allele specific enhancers with either high or low RNAPII signal.**

**ATAC peaks downstream of allele specific enhancers with high RNAPII signal**

| Peaks Away | ATAC Q-Value | K27ac Q-Value | K4me1 Q-Value | Distance |
|---|---|---|---|---|
| 1 | 4.1E-26 | 1.22E-55 | 2.81E-72 | 3174.132 |
| 2 | 7.11E-18 | 7.15E-35 | 3.26E-40 | 10477.03 |
| 3 | 2.87E-14 | 2.49E-27 | 1.07E-30 | 18833.72 |
| 4 | 4.05E-10 | 2.36E-17 | 7.17E-21 | 27610.45 |
| 5 | 0.000324 | 2.34E-10 | 1.68E-13 | 35975.81 |
| 6 | 0.000371 | 1.45E-08 | 1.19E-12 | 44844.17 |
| 7 | 0.000272 | 1.3E-05 | 1.68E-13 | 54473.27 |
| 8 | 0.037131 | 0.00033 | 1.3E-08 | 63664.44 |
| 9 | 0.008063 | 0.000174 | 2.77E-07 | 73470.75 |
| 10 | 0.152029 | 0.002843 | 9.68E-06 | 82895.08 |
| 11 | 2.182492 | 0.030789 | 4.43E-05 | 92433.13 |
| 12 | 33.72391 | 0.144833 | 0.001272 | 102237.2 |
| 13 | 4.803357 | 0.098187 | 0.001081 | 112580.3 |
| 14 | 1.210011 | 0.035371 | 0.000421 | 121988.6 |
| 15 | 8.097056 | 0.295751 | 0.004072 | 131566.5 |
| 16 | 3.601671 | 0.643326 | 0.008129 | 141130.1 |
| 17 | 16.38582 | 1.52541 | 0.080605 | 150639 |
| 18 | 35.21653 | 1.219326 | 0.081741 | 160408.1 |
| 19 | 36.14848 | 7.607946 | 0.696776 | 170048.9 |
| 20 | 72.94739 | 14.12048 | 1.987121 | 179581.8 |
| 21 | 74.34156 | 39.06271 | 0.914002 | 188769.9 |
| 22 | 374.9811 | 63.82918 | 19.42994 | 199011.5 |
| 23 | 90.60817 | 22.06589 | 38.8355 | 209275.1 |
| 24 | 362.827 | 190.8466 | 34.12068 | 219282.9 |
| 25 | 273.8584 | 302.3124 | 88.28709 | 229911.1 |
| 26 | 81.56936 | 314.3522 | 37.66934 | 239714.2 |
| 27 | 135.1474 | 151.8752 | 72.38432 | 249539.2 |
| 28 | 560.4686 | 352.8446 | 277.0628 | 259076.6 |
| 29 | 578.6429 | 542.8033 | 315.2243 | 268837.4 |
| 30 | 893.5414 | 616.0511 | 269.676 | 278526.4 |
| 31 | 1052.13 | 578.4087 | 692.8376 | 288087.6 |
| 32 | 263.8834 | 975.0438 | 391.2057 | 298734.4 |
| 33 | 588.0324 | 1435.581 | 528.1608 | 308241.9 |
| 34 | 1422.247 | 1407.344 | 564.5849 | 318234 |
| 35 | 489.6612 | 1237.854 | 131.9067 | 328016.1 |
| 36 | 540.0336 | 1241.829 | 182.5695 | 338120.5 |
| 37 | 531.7491 | 1447.54 | 85.59545 | 347976.5 |
| 38 | 326.6261 | 744.7664 | 59.36504 | 357786.4 |
| 39 | 104.7225 | 413.6851 | 203.0817 | 367863.9 |
| 40 | 426.9381 | 441.1174 | 231.5403 | 377937.8 |

**ATAC peaks upstream of allele specific enhancers with high RNAPII signal**

| Peaks Away | ATAC Q-Value | K27ac Q-Value | K4me1 Q-Value | Distance |
|---|---|---|---|---|
| 1 | 1.22E-25 | 2.28E-61 | 5.24E-63 | 3443.309 |
| 2 | 5.69E-17 | 2.65E-38 | 2.03E-42 | 11143.11 |
| 3 | 9.97E-11 | 4.81E-27 | 6.05E-31 | 19367.46 |
| 4 | 5.88E-07 | 1.52E-17 | 6.97E-21 | 28046.23 |
| 5 | 1.96E-05 | 2.46E-11 | 1.41E-15 | 37028.94 |
| 6 | 3.84E-05 | 1.01E-08 | 8.32E-13 | 46705.81 |
| 7 | 0.000326 | 8.97E-09 | 6.28E-12 | 56230.83 |
| 8 | 0.003793 | 4.3E-06 | 6.83E-09 | 65735.41 |
| 9 | 0.005269 | 0.002356 | 1.21E-08 | 74829.94 |
| 10 | 0.048998 | 0.001527 | 1.14E-05 | 84410.74 |
| 11 | 1.098134 | 0.000449 | 0.000391 | 93901.26 |
| 12 | 5.595652 | 0.015113 | 0.001441 | 103979.9 |
| 13 | 36.2085 | 1.399854 | 0.016608 | 114209 |
| 14 | 29.40115 | 0.824644 | 0.024789 | 123958.9 |
| 15 | 15.12526 | 12.88933 | 0.025318 | 133829.8 |
| 16 | 15.02258 | 1.724285 | 0.03715 | 143270.5 |
| 17 | 5.821238 | 2.289546 | 0.339104 | 153076.1 |
| 18 | 46.82668 | 0.964982 | 0.764203 | 162660.8 |
| 19 | 58.00963 | 0.827566 | 0.0906 | 172446.3 |
| 20 | 35.69868 | 4.259461 | 0.149858 | 182431.1 |
| 21 | 37.48786 | 5.0437 | 1.26923 | 192283.2 |
| 22 | 28.32624 | 13.79159 | 2.24888 | 201737.4 |
| 23 | 63.76953 | 32.01725 | 12.8526 | 211415.4 |
| 24 | 155.2147 | 75.38659 | 24.24967 | 221307.5 |
| 25 | 153.352 | 253.1979 | 32.00922 | 230781.5 |
| 26 | 286.0938 | 241.5021 | 31.62523 | 240467.4 |
| 27 | 200.8416 | 195.8677 | 71.7323 | 250452.4 |
| 28 | 468.8489 | 376.3457 | 16.15865 | 259794.6 |
| 29 | 188.6159 | 345.6827 | 44.00091 | 269882.6 |
| 30 | 562.8537 | 552.6525 | 68.65264 | 280430.4 |
| 31 | 597.0781 | 476.1396 | 127.8117 | 290106 |
| 32 | 920.8624 | 741.9614 | 489.2754 | 299887.7 |
| 33 | 1602.441 | 1008.767 | 940.8364 | 309542.6 |
| 34 | 1284.395 | 916.7598 | 889.7291 | 318757.2 |
| 35 | 433.2802 | 1512.148 | 344.3508 | 328299.2 |
| 36 | 620.1687 | 1417.997 | 460.1093 | 337660.8 |
| 37 | 1195.916 | 1762.889 | 902.4615 | 346802.8 |
| 38 | 1398.564 | 1908.637 | 1582.69 | 355723.6 |
| 39 | 1655.878 | 1733.448 | 1322.98 | 364894.1 |
| 40 | 792.1868 | 982.6174 | 1576.65 | 374117.5 |

**ATAC peaks downstream of allele specific enhancers with low RNAPII signal**

| Peaks Away | ATAC Q-Value | K27ac Q-Value | K4me1 Q-Value | Distance |
|---|---|---|---|---|
| 1 | 1.43E-11 | 5.31E-20 | 7.29E-35 | 6864.823 |
| 2 | 5.47E-06 | 4.64E-10 | 6.11E-16 | 21901.92 |
| 3 | 1.41E-01 | 1.58E-02 | 1.99E-07 | 36557.3 |
| 4 | 1.15E+00 | 1.59E-01 | 1.30E-05 | 52347 |
| 5 | 3.63E+00 | 1.78E-01 | 1.67E-04 | 67663.93 |
| 6 | 5.13E+00 | 9.51E-02 | 1.20E-03 | 83154.93 |
| 7 | 1.96E+00 | 1.11E+00 | 2.05E-02 | 98043.94 |
| 8 | 6.27E+00 | 1.05E+01 | 8.34E-02 | 112363.8 |
| 9 | 3.16E+01 | 1.02E+01 | 4.49E-01 | 127038.6 |
| 10 | 2.22E+01 | 5.51E+00 | 7.92E-02 | 141746 |
| 11 | 2.07E+02 | 2.44E+01 | 2.78E-01 | 156260.4 |
| 12 | 2.12E+02 | 2.77E+01 | 7.86E-01 | 171143.2 |
| 13 | 2.34E+02 | 1.82E+02 | 1.93E+01 | 186045.8 |
| 14 | 1.13E+01 | 8.32E+01 | 1.34E+00 | 201559.2 |
| 15 | 1.35E+02 | 2.44E+02 | 9.53E+00 | 215906.7 |
| 16 | 7.90E+02 | 3.15E+02 | 2.67E+01 | 231010.4 |
| 17 | 5.73E+02 | 3.96E+02 | 1.04E+01 | 245186 |
| 18 | 5.29E+02 | 2.08E+02 | 4.87E+01 | 259900 |
| 19 | 4.34E+02 | 2.51E+02 | 5.75E+01 | 274520.7 |
| 20 | 9.25E+01 | 3.60E+01 | 3.75E+01 | 289699.8 |
| 21 | 1.61E+02 | 3.83E+01 | 1.16E+02 | 304565.9 |
| 22 | 9.58E+01 | 4.67E+01 | 1.64E+02 | 318760.7 |
| 23 | 1.02E+02 | 1.04E+02 | 6.87E+01 | 332972.7 |
| 24 | 3.22E+02 | 9.30E+02 | 1.70E+02 | 346848.9 |
| 25 | 1.61E+02 | 5.09E+02 | 2.71E+02 | 360565.8 |
| 26 | 2.54E+02 | 1.25E+03 | 1.91E+02 | 374257.5 |
| 27 | 5.14E+02 | 1.01E+03 | 2.26E+02 | 387881.3 |
| 28 | 6.04E+02 | 8.61E+02 | 2.16E+02 | 401403.2 |
| 29 | 5.50E+02 | 9.38E+02 | 1.61E+02 | 415481.1 |
| 30 | 9.19E+02 | 1.78E+03 | 2.59E+02 | 428959.5 |
| 31 | 8.28E+02 | 1.93E+03 | 3.44E+02 | 443123.2 |
| 32 | 8.63E+02 | 1.45E+03 | 5.57E+02 | 457244 |
| 33 | 9.16E+02 | 6.58E+02 | 4.30E+02 | 471535.7 |
| 34 | 6.70E+02 | 6.35E+02 | 2.43E+02 | 484933.6 |
| 35 | 3.50E+02 | 5.29E+02 | 2.37E+02 | 498531.7 |
| 36 | 8.32E+02 | 1.21E+03 | 1.51E+02 | 512305.9 |
| 37 | 4.06E+02 | 6.29E+02 | 2.10E+02 | 526051.7 |
| 38 | 2.11E+02 | 2.65E+02 | 1.75E+02 | 539217 |
| 39 | 5.99E+01 | 1.20E+02 | 1.54E+02 | 552646.1 |
| 40 | 1.60E+02 | 2.28E+02 | 1.91E+02 | 565543.2 |

**ATAC peaks upstream of allele specific enhancers with low RNAPII signal**

| Peaks Away | ATAC Q-Value | K27ac Q-Value | K4me1 Q-Value | Distance |
|---|---|---|---|---|
| 1 | 2.71E-13 | 5.61E-19 | 1.31E-38 | 7087.422 |
| 2 | 3.98E-07 | 2.83E-13 | 7.64E-21 | 21522.51 |
| 3 | 2.58E-03 | 1.21E-08 | 1.63E-12 | 36222.96 |
| 4 | 1.19E-03 | 1.40E-06 | 1.82E-08 | 51012.07 |
| 5 | 3.22E-02 | 4.96E-04 | 6.66E-06 | 65668.07 |
| 6 | 2.01E+00 | 2.71E-02 | 2.26E-03 | 80867.83 |
| 7 | 6.22E+00 | 8.90E-01 | 1.81E-02 | 96694.67 |
| 8 | 7.99E+00 | 2.39E-01 | 1.21E-01 | 111059.7 |
| 9 | 1.35E+01 | 5.46E-01 | 3.75E-01 | 124678.3 |
| 10 | 1.80E+01 | 9.50E-01 | 1.64E-01 | 139489.9 |
| 11 | 1.15E+01 | 3.22E+00 | 8.63E-02 | 153732.4 |
| 12 | 3.13E+02 | 1.62E+01 | 3.38E+00 | 168148.1 |
| 13 | 3.58E+02 | 1.74E+01 | 1.44E+01 | 182627.4 |
| 14 | 2.61E+01 | 4.34E+00 | 3.26E+00 | 197568.4 |
| 15 | 6.24E+01 | 1.29E+01 | 2.44E+00 | 212060.8 |
| 16 | 3.27E+02 | 1.50E+02 | 1.59E+01 | 226647.4 |
| 17 | 6.74E+02 | 3.93E+02 | 8.03E+01 | 241845 |
| 18 | 7.58E+02 | 9.60E+02 | 1.06E+02 | 255448.4 |
| 19 | 7.71E+02 | 1.03E+03 | 6.62E+01 | 270134 |
| 20 | 4.70E+02 | 8.79E+02 | 7.44E+01 | 284663.9 |
| 21 | 1.33E+03 | 7.76E+02 | 4.00E+02 | 298276 |
| 22 | 7.15E+02 | 4.44E+02 | 2.81E+02 | 312636.9 |
| 23 | 1.43E+03 | 7.29E+02 | 3.65E+02 | 326230.2 |
| 24 | 6.01E+02 | 7.03E+02 | 2.24E+02 | 340447 |
| 25 | 4.49E+02 | 5.03E+02 | 1.43E+02 | 354310.8 |
| 26 | 4.13E+01 | 1.28E+02 | 2.67E+01 | 367892.4 |
| 27 | 2.37E+02 | 1.94E+02 | 5.35E+01 | 381293.9 |
| 28 | 9.71E+02 | 8.49E+02 | 2.44E+02 | 394450.9 |
| 29 | 1.36E+03 | 1.82E+03 | 8.13E+02 | 408076.4 |
| 30 | 1.54E+03 | 1.68E+03 | 6.06E+02 | 421415.3 |
| 31 | 1.04E+03 | 1.91E+03 | 6.35E+02 | 435504.1 |
| 32 | 8.69E+02 | 1.81E+03 | 6.50E+02 | 448501.7 |
| 33 | 4.06E+02 | 1.63E+03 | 2.88E+02 | 461196.3 |
| 34 | 4.19E+02 | 1.26E+03 | 3.27E+02 | 474211.3 |
| 35 | 3.80E+02 | 1.04E+03 | 1.68E+02 | 487094.7 |
| 36 | 3.58E+02 | 6.59E+02 | 9.46E+01 | 499704.5 |
| 37 | 4.73E+02 | 7.88E+02 | 1.06E+02 | 512721.8 |
| 38 | 3.25E+02 | 8.09E+02 | 2.04E+02 | 526057.4 |
| 39 | 1.51E+02 | 2.21E+02 | 4.01E+02 | 539180.2 |
| 40 | 6.10E+02 | 6.00E+02 | 4.61E+02 | 551648.4 |

Allele specific enhancers were split up into those with greater than 500 reads per million of RNAPII signal and those with less. For each ATAC peak upstream and downstream of allele specific enhancers high or low RNAPII signal, we calculated the adjusted P-Value (Q-Value) of 129 and CAST read counts for ATAC, H3K27ac, and H3K4me1 data. The last column is the peaks distance from the allele specific enhancer.
